# Extracellular vesicles regulate metastable phenotypes of lymphangioleiomyomatosis cells via shuttling ATP synthesis to pseudopodia and activation of integrin adhesion complexes

**DOI:** 10.1101/2024.09.09.611297

**Authors:** Anil Kumar Kalvala, Ashok Silwal, Bhaumik Patel, Apoorva Kasetti, Kirti Shetty, Jung-Hung Cho, Gerard Lara, Beth Daugherity, Remi Diesler, Venkatesh Pooladanda, Bo R Rueda, Elizabeth Petri Henske, Jane J. Yu, Maciej Markiewski, Magdalena Karbowniczek

## Abstract

Pulmonary lymphangioleiomyomatosis (LAM) is metastatic sarcoma but mechanisms regulating LAM metastasis are unknown. Extracellular vesicle (EV) regulate cancer metastasis but their roles in LAM have not yet been investigated. Here, we report that EV biogenesis is increased in LAM and LAM EV cargo is enriched with lung tropic integrins, metalloproteinases, and cancer stem cell markers. LAM-EV increase LAM cell migration and invasion via the ITGα6/β1-c-Src-FAK-AKT axis. Metastable (hybrid) phenotypes of LAM metastasizing cells, pivotal for metastasis, are regulated by EV from primary tumor or metastasizing LAM cells via shuttling ATP synthesis to cell pseudopodia or activation of integrin adhesion complex, respectively. In mouse models of LAM, LAM-EV increase lung metastatic burden through mechanisms involving lung extracellular matrix remodeling. Collectively, these data provide evidence for the role of EV in promoting LAM lung metastasis and identify novel EV-dependent mechanisms regulating metastable phenotypes of tumor cells. Clinical impact of research is that it establishes LAM pathway as novel target for LAM therapy.

## Introduction

Pulmonary lymphangioleiomyomatosis (LAM) is low-grade, understudied, and metastasizing sarcoma, predominately affecting woman, and manifesting as proliferation of tumor smooth muscle-like cells within the lungs, which ultimately leads to lung damage and failure^1–4^. It develops in association with tuberous sclerosis complex (TSC) or as a sporadic form. Both TSC and sporadic LAM result from germline or somatic *TSC1/TSC2* mutations, respectivly^5,6^ that prevent the inhibition of the mechanistic target of rapamycin (mTOR) by TSC1/TSC2 complex^7,8^. The metastatic potential of LAM, which we discovered, and the origin, possibly, from uterus, renal angiomyolipoma, or from unknown site are now well-accepted^1–4^. However, mechanisms regulating LAM metastasis are unknown.

Here we report on a role of extracellular vesicles (EV) in the promotion of LAM metastasis. EV, which are released from cells, including cancer cells, to mediate cell-to-cell communication through their cargo, and promote metastasis. We previously reported that EV derived from *Tsc1*–null neuronal progenitors block differentiation of recipient wild-type progenitors via the activation of Notch1/mTOR pathways, phenocopying *Tsc1*-null cells, and that mTORC1 hyperactive LAM surrogate cells secrete EV, thereby affecting target cells via activation of Notch1/mTOR^9^. Consistently, LAM surrogate cells have increased EV biogenesis and cargo that enhance VEGF secretion and viability of recipient fibroblast compared to TSC2 addback cells^10^. Therefore, we hypothesize that EV contributes to LAM metastasis, especially in the context of the abundance of EV in lung microenvironment. The lung endothelium, alveolar macrophages, fibroblasts, and epithelial cells are the sources of EV that contribute to asthma, chronic obstructive pulmonary disease (COPD), pulmonary hypertension, and lung cancer^11–14^.

EV biogenesis and uptake are regulated by the classical and non-classical endocytic pathways ^15–23^. EV from epithelial cancers (carcinomas) and epithelial cancer stem cells (CSCs) or from the tumor microenvironment (TME) influence CSCs, premetastatic niche, metastasis, and response to therapy^24–29^. These EV transport growth factors, integrins, non-receptor tyrosine kinase protooncogene c-Src, and focal adhesion kinase (FAK), and can regulate angiogenesis, vascular permeability, premetastatic niche, and seeding of target organs by tumor cells^24–27,30–34^. EV-derived integrins (EV-ITGs) regulate anchorage-independent (i.e. in the circulation) growth of tumor cells and their organotropism^33,35^. The lung-tropic EV-ITGs: ITGα6, ITGβ4, and ITGβ1^35^, bind to the lung-resident fibroblasts and epithelial cells to promote lung metastasis via the induction of *S100*^35^. S100s promote cancer progression by altering the premetastatic niche and cancer cells^36–41^. Despite to advancement in expanding roles of EV in carcinomas, their functions in non-epithelial malignancies, especially sarcomas, including LAM, are understudied. Limited evidence defines the potential roles for EV in regulating tumor angiogenesis, adhesion, and migration of non-epithelial/mesenchymal malignant cells^28,29,42–44^.

Cancer cell plasticity is linked to stemness, anoikis resistance, and increased metastatic potential^45–54^. Carcinoma cells oscillate between a proliferative/differentiated and invasive/dedifferentiated phenotype (metastable/hybrid phenotypes)^47–54^. Cancer cell plasticity and hybrid metastable phenotypes are also observed in non-epithelial tumors^45,55–60^. Sarcoma CSCs regardless of origin form clusters or “sarcospheres” in the circulation^3,60–63^ and share stem cell characteristics such as nestin and CD44 expression and high levels of active aldehyde dehydrogenase (ALDH)^60,61^. CD44 associates with metastable phenotypes of mesenchymal tumors^55^. LAM cells express several CSC markers including CD44, ITGs and ALDH^3,4,62–65^ and “stem-like state” LAM cells’ subpopulation exists^66^. Therefore, we theorize that EV promote LAM metastasis through the regulation of LAM cell plasticity and metastable phenotypes.

## Materials and Methods

### Cells

<u>ELT3</u>: Tsc2-null uterine leiomyoma-derived from the Eker rat model of TSC, by C. Walker^67,68^. <u>621-101</u>: human LAM surrogate cells (LAM-associated angiomyolipoma-derived) with bi-allelic *TSC2* mutations^69,70^ (from Drs. Henske and Yu); <u>621-103</u>: TSC2-reexpressing 621-101 cells (from Drs. Henske and Yu); <u>621L9</u>: 621-101 cells stably expressing luciferase (from Dr. Yu)^71^. The cell number and viability were determined before plating. 621-101 and 621-103 were cultured in standard Dulbecco’s modified Eagle’s medium (DMEM) with the addition of 10% Fetal Bovine Serum (Corning, #35-010-CV), 1x penicillin/streptomycin (Corning, #30-002-CL), and 5 ug/ml plasmocin prophylactic (Invivogen, San Diego, CA). Plates were incubated at 37°C with 5% CO2 until cells were approximately 80% confluent. For experiments cells were plated at equal numbers in DMEM medium containing 10% FBS depleted of EV by standard ultracentrifugation^9,72^. To generate spheroids, cells were seeded on ultra-low attachment plates at density of 6000 cells/mL unless otherwise specified^73^. Briefly, cells were cultured in DMEM/F-12 (Corning, #10-090-CV) with the addition of 3% EV free FBS (Corning, #35-010-CV), 1x non-essential amino acids (Corning, #25-025-Cl), 1x penicillin/streptomycin (Corning, #30-002-CL), 1x N2 supplement (Gibco; #17502-048), 1x B27 without Vit A (Gibco, #12587-010), 20 ng/ml EGF (PROSPEC, #cyt-217), 20 ng/ml FGF (PROSPEC, #cyt-218), 10 ng/ml LIF (Peprotech, #300-05), 100 µM β-mercaptoethanol (Gibco, #21985-023) (sphere media). The E15.5 embryo neural tube (NT) derived cells were cultured in 15% EV-free FBS DMEM/F12 media supplemented with 20 ng/ml EGF, 20 ng/ml bFGF, 20 ng/ml IGF, 1% B-27, 1% N2 supplement, 1% penicillin/streptomycin, as described^9,74^.

### Transfection to generate stable cell lines

621-101 and TSC2 addback cells were infected by lentiviral transduction with dual-color fluorescent reporter for CD63-positive exosome secretion and uptake (Addgene plasmid # 172118), as previously described^9^. pLenti-pHluorin_M153R-CD63-mScarlet was a gift from Alissa Weaver (Addgene plasmid # 172118; http://n2t.net/addgene:172118; RRID: Addgene_172118)^75^.

### EV isolation

#### Ultrafiltration followed by size exclusion chromatography (SEC) method

2D culture system was used to mimic primary tumor-like conditions. Whereas, 3D culture system using ultra-low attachment plates was used to mimic metastasizing tumor-like conditions (i.e. circulation). For isolation of primary tumor-like EV cells were plated in 150 mm dish at the seeding density of 5×10^6^ cells in DMEM with 10% EV free FBS and allowed to incubate for 72 hours. For isolation of metastasizing tumor-like EV cells were grown as spheres. Conditioned media was subjected to serial centrifugation steps at 500 x g (5 minutes), 2000 x g (10 minutes), and 10,000 x g (30 minutes) to remove all cell debris. Then, supernatant was filtered through 0.22 µm syringe filters and passed through pre-equilibrated Amicon 100 kDa ultrafilters (Millipore Sigma, #UFC910024) using three consecutive centrifugations for 30 minutes at 3000 x g^76^. Next, concentrated EV were eluted with PBS and 100 µl of this concentrate was passed through Size exclusion chromatography (SEC) columns (Cell guidance systems, #Ex03). Columns were washed several times with PBS. Six fractions were collected. The first fraction represented EV-depleted media (EDM), while the second fraction represented EV.

#### Ultracentrifugation combined with sucrose cushion method (from conditioned media or plasma)

EV were purified by initial centrifugation followed by filtration (0.22 µm), standard ultracentrifugation with or without EV pelleting in density 30% sucrose gradient^72,77–79^. Equal volumes of diluted plasma (≥ 200 µL diluted in 4 mL of PBS) were used for EV isolation^72^.

### EV characterization

Equal quantities of initial bio-fluid, initial number of plated cells, or time of conditioning were used. The fluorescence-activated cell sorting (FACS), dynamic light scattering (DLS), and nanoparticle tracking analysis (NTA) were used^9,72^. Samples were sent to the Texas Tech University College of Arts and Sciences for transmission electron microcopy (TEM) analyses and lipid bilayer detection. Western immunoblotting assessed the presence of proteins. The EV samples and bio-fluid after EV depletion was loaded at equal quantities per *JEV*^72^. EV were examined for transmembrane, cytosolic (GAPDH), and non-EV proteins (albumin), and other organelles: nucleus, mitochondria, ER, and Golgi. The EV used in studies were normalized by total amount of protein in the sample. For assays, equal quantities of EV and *JEV* controls were used unless otherwise specified^72^.

#### EV Characterization by FACS

Isolated exosomes were characterized as previously described^9,80^. Briefly, aldehyde/sulfate beads (Interfacial Dynamics, Grand Island, NY, USA) were incubated with capture human CD63 (BD Biosciences), human CD9 (Biolegend), and mouse CD9 (BioLegend, San Diego, CA, USA) antibodies and then with mouse plasma or conditioned media. EV-coated beads were incubated with Human CD63 (Biolegend), Human CD9 (Biolegend), Mouse CD63 (Biolegend), and Mouse CD9 (BD Biosciences) antibodies and analyzed by FACS.

#### EV Characterization by NTA

Isolated EV were analyzed by NTA (System Biosciences, version 2.3 build 2.3.5.0033.7-Beta7 of the NTA software). The EV size and particle number were evaluated.

#### EV Characterization by DLS

using dynamic light scattering (Malvern Zetasizer ultra red, # ZSU3305), zeta size and particle distribution were evaluated. The particle distribution was reported in percent intensity defined as a plot of the relative percentage of particles contained in various size classes based upon the intensity of scattered light.

#### EV Characterization by TEM

Briefly, EVs were washed with 0.05 M Cacodylate buffer (3x) and post-fixed with 1% osmium tetroxide for 1 hour, followed by washing (3x). EVs were dehydrated through increasing ethanol concentrations (25% to 100%) and acetone (100%), then infiltrated with plastic (4:1, 1:1, 1:4 acetone) and embedded in Epon for 48 hours. Blocks were trimmed and mounted in a microtome to cut 1 µm thick sections, which were stained with methylene blue azure II, covered with permount, and examined under a microscope. For thin sectioning, blocks were re-trimmed, cut to 70-90 nm with a diamond knife, and placed on copper grids. Grids were stained with 1% uranyl acetate solution from a 4% stock using NERL water, washed, dried, and imaged using a Hitachi H-7650 TEM^81^.

### EV labeling, uptake, and biogenesis

Isolated EV were labeled using ExoGlow (SBI, EXOGP300A-1 or EXOGM600A-1) according to manufacturer and approximately 100 µg of EV solution was added to the cells for 6 hours. Next, cells were trypsinized and subjected for EV uptake analysis using FACS. For inhibition of EV uptake, 10 µM Dyngo4a was added 3 hours prior to EV treatment.

For inhibition of EV biogenesis cells were treated with 0.1 µM Tipifarnib or 10 µM GW4869, and media was subjected for EV isolation and characterization by FACS^15,20–23,82^.

### Inhibitors for studies using adherent cells

Bosutinib (1 µM), GW4869 (10 µM), Tipifarnib (0.1 µM) were added to 621-101 cells and maintained for 24 hours. Next, cells were trypsinized and plated to transwell inserts in the presence of inhibitors and EV. Dyngo4a (10 µM) was added to 621-101 cells 3 hours prior to EV exposure. Cells were then incubated for 16 and 72 hours for migration and invasion, respectively.

### Inhibitors for studies using spheres

Bosutinib (1 µM), GW4869 (10 µM), Tipifarnib (0.1 µM) and Dyngo4a (10 µM) were added to day 0 and day 2 621-101 spheres, and spheres were allowed to grow for 7 days. Next, D7 spheres were subjected to downstream assays.

### Cell lysis

Cells were washed with ice-cold PBS and lysed on ice for 15-20 minutes with RIPA buffer supplemented with PhosSTOP^TM^ (Roche, 4906845001) and protease inhibitors (Thermo scientific, A32965) for whole cell lysates (WCL). WCL were cleared by the centrifugation at 14,000 RPM for 15 min at 4^0^C and protein concentration was determined using the Bradford assay (Bio-Rad Laboratories, 5000006).

### Western immunoblotting

Protein lysates were boiled for 10 min and subjected to SDS-PAGE electrophoresis using 4%-12% precast gels (Invitrogen, NP0336BOX, and NP0322BOX). Primary antibody binding was detected using HRP-conjugated anti-mouse or anti-rabbit antibody (Invitrogen) and chemiluminescence (Thermo Scientific). Primary antibodies were used at a dilution of 1:1,000 in 5% BSA/TBST solution, and secondary antibodies at 1:10,000 in 5% milk/TBST unless otherwise specified (Tab. 1).

### Quantitative (q) real time (RT)-PCR

RNA was extracted using Rneasy plus mini kit (Qiagen) and cDNA was generated using High-Capacity RNA-to-cDNA^TM^ kit (Applied Biosystems). The qRT-PCR was performed using High Capacity cDNA Synthesis Kit, Fast SybrGreen and StepOne Plus (Applied Biosystems). The comparative Ct method (2-ΔΔCt) and RT2 profiler PCR Array Data Analysis (SAB Biosciences) was used to determine fold differences between the target gene and the housekeeping gene GAPDH. Primer sequences were established based on https://pga.mgh.harvard.edu/primerbank/ (Tab. 2).

### Immunofluorescence and confocal microscopy

#### For adherent cells

621-101 cells cultured overnight on coverslips and fixed with 4% formaldehyde in PBS for 15 minutes at room temperature, rinsed with PBS, and then exposed to blocking buffer (5% BSA/0.3% Triton-X-100 in PBS) for 1 hour at room temperature. This was followed by 1-hour incubation with F-Actin (1:2000, Spirochrome, #SPY555-actin, Tab. 1) followed by overnight incubation with anti-Rabbit p-paxillin (Y118) (1:50, Cell Signaling Technology, #2541S, Tab.1). The next day, the cells were rinsed with PBS and incubated with anti-Rabbit-FITC (1:400 dilution) for 1 hour at room temperature and rinsed with PBS. Cells were mounted with ProLong™ Diamond Antifade Mountant (Thermofischer scientific, #P36965) and imaged using Nikon AX R confocal microscope.

#### For sphere cells

621-101 spheres were plated onto collagen-coated coverslips and allowed to migrate for 6 hours before staining with TFAM and p-paxillin (Y118) antibodies, and WGA (Tab.1). Fluorescence was observed with Nikon AX R confocal microscope and quantified using Nikon Elements Advanced Research Image-Analysis software. Data is expressed as mean fluorescence intensity (MFI) or Pearson correlation coefficient for colocalization.

### Immunohistochemistry

Sections were deparaffinized, incubated with primary antibody, S100A4 (1:800, Rabbit mAb, #13018, Cell Signaling Technology) and biotinylated secondary antibodies-Rabbit specific HRP/DAB (ABC) Detection IHC kit (#PK-4000, Vector Laboratories, Inc, CA, USA).

### Trichome Mason staining

Masson’s Trichrome staining was conducted as outlined by Burkholder (1974). FFPE tissue sections were deparaffinized, rehydrated, stained with Carazzi hematoxylin, followed by 1% Briebrich scarlet and 1% acid fuchsin. After decolorizing with 1% phosphotungstic acid, the sections were stained with 1% light green, rinsed in ethanol and xylene, and coverslipped.

### Scratch migration assay

621-101 cells were seeded at a density of 1.2 million cells per well in EV-free DMEM/F12 complete media on a 6-well plate. The following day, a scratch was created vertically down the center of each well using a commercially available scratcher and images were captured every 2 hours using a Citation 5. The wound healing efficiency was determined from three selected fields at each time point by calculating the difference between the original wound area and the post-migration area, divided by the original wound area.

### Transwell migration and invasion assays

Transwell chambers were coated with a 100 µl solution containing 50 µg/cm^2^ rat tail collagen IV (for migration) or 300 µg/ml growth factor reduced Matrigel matrix (for invasion), followed by a 2-hour incubation at room temperature with gentle shaking, or at 37°C in a CO2 incubator, respectively. The 0.5 ml volume of single-cell suspensions (25000-50000 cells/well) in serum-free medium were plated into 24-well inserts. The 0.75 ml of 10% FBS complete EV free DMEM/F12 media was added to companion plate wells. Chambers were incubated for 16 hours (migration) or 72 hours (invasion) at 37°C with 5% CO2. Non-migrated/invaded cells were removed from the upper chamber using cotton swabs, and remaining cells at the bottom were stained, air-dried, scanned using Aperio, and quantified using ImageJ software (Imagej 1.53k, NIH, USA).

### Primary and secondary sphere assays

Cells were seeded at density of 500 cells per well into ultra-low attachment round or flat bottom 96-well plates in sphere media. Plates were centrifuged daily at 200 x g for 5 minutes. Cells were treated with EV or inhibitors on day 0 and day 2 and grown for 7 days or as indicated. The sphere size was determined using either confocal microscopy or Citation 5. Secondary sphere formation assay was performed with slight modifications of previous protocol^83^. Briefly, on day 7, primary spheres were dissociated with trypsin, neutralized with serum media, and filtered through a 70 µm nylon mesh to form a single-cell suspension. Cell count and viability were assessed with a cell counter. Next, 500 cells per well were seeded into low-attachment 96-well plates containing sphere media. After seven days, spheres were imaged and sphere size was assessed using NIKON confocal microscope and NIS-Elements AR 5.42.03 64-bit software respectively.

### Sphere cell proliferation

Single primary spheres were labeled with EdU labeling solution at day 6 and final concentration of 10 µM of Click-iT^TM^ EdU Alexa Fluor^TM^ 488 (Invitrogen, #C10337) in sphere media, incubated for 4 hours, then transferred to 1.5 ml tubes, washed once with 3% BSA in PBS and fixed with 3.7% formaldehyde in PBS for 30 minutes. Subsequently, spheres were washed twice with 3% BSA in PBS and permeabilized with 0.5% Triton X-100 in PBS for 30 minutes at room temperature followed by three washes with 3% BSA in PBS. The spheres were incubated in Click-iT reaction cocktail in the dark for 30 minutes at room temperature, washed twice with 3% BSA in PBS, counterstained with Fluoro-Gel II containing DAPI (Electron Microscopy Sciences, #17985-51) for 45 minutes, washed once with 3% BSA in PBS before mounting in ProLong™ Diamond Antifade Mountant (Invitrogen, Catalog no P36961). Images were captured using Nikon AX-R confocal Microscope at 20x magnification, with a zoom size of 4 and 1-µm-thick Z-stacks spanning the entire sphere. The images were analyzed on ImageJ using the ‘Multi-point’ tool to count the percentage of EdU positive cells against the nuclear counterstain in the same sphere region.

### ALDH assay

The ALDH activity was measured using ALDEFLUOR^TM^ Kit (StemCell Technologies, Canada).

### Sphere migration assay

621-101 cells were cultured in sphere-forming media and treated with EVs or inhibitors on days 0 and 2. On day 7, spheres were transferred to collagen-coated 96-well plates (50 µg/cm²) and allowed to settle for 2 hours, next imaged using Citation 5, then cells were allowed to migrate in a CO₂ incubator for 24 hours imaged again. Migration was assessed by drawing 10 lines from the sphere edge to the furthest point of migrated cells using Citation 5 to quantify migrated distance. Data were analyzed with GraphPad Prism.

### Time-lapse imaging of sphere cell migration and trajectory plots generation

Day 7 spheres were transferred to a flat-bottom 96-well plate coated with 50 ug/cm^2^ of rat tail Type-I collagen (Advanced BioMatrix, #5153-100MG) allowed to settle at the bottoms for 2 hours in a CO2 incubator before time-lapse imaging using Nikon Microscope AX-R and 4x objective. Spheres were imaged every 10 minutes over 12-16 hours. The captured image had 30 pixels corresponding to 100 µm. Next, migrated cells were tracked using the ImageJ Manual Tracking plugin to determine their positional values (x, y) at each time point. The output from the Manual Tracking plugin was further processed using the Ibidi Chemotaxis and Migration Tool V2.0 to generate trajectory plots of migrating cells and determine their distance and velocity.

### Measurement of ATP level

Cellular ATP and lactate levels were assessed using the Enzylight ATP Assay Kit (BioAssay Systems, USA, #EATP-100) following the manufacturer’s protocols. 621-101 cells, cultured as spheres for seven days, were lysed with 50 µl of PBS. ATP was determined by the amount of light emitted after the reaction of D-luciferin and ATP catalyzed by luciferase. The luminescent signal was recorded using the luminometry mode of a plate reader (BioTek Cytation5). ATP levels were normalized to the protein content and reported as µM per mg of protein.

### Cell body and pseudopodia isolation

Pseudopods were obtained from cell bodies as described previously^84^. In brief, cell culture inserts with 3.0 µm-pore polycarbonate membranes (CELLTREAT, #230609) were coated with collagen at a concentration of 50 µg/cm^2^ for 2 hours at room temperature, rinsed with PBS, and seeded with cells from dissociated spheres. Cells were allowed to migrate for 6 hours. The pseudopods were collected by gently scraping the top surface of the insert with a cotton swab and transferring them into lysis buffer. To isolate the cell bodies, the undersides of the inserts were scraped to remove the pseudopods, and the remaining cell bodies were collected in PBS for subsequent analysis via immunoblotting and ATP assay.

### Integrin adhesion complexes isolation

The isolation of integrin-associated adhesion complexes was carried as described previously^85^ with minor modifications. Spheres were transferred to a collagen-coated plates for 6 hours. Next, plates were washed twice with pre-warmed DMEM-HEPES to remove non-adherent cells, followed by an 8-minute incubation with a 6 mM solution of DTBP cross-linker (Thermo Fisher Scientific) in DMEM-HEPES at 37°C. The cross-linker was quenched with 150 µl of 1M Tris-HCl, pH 8. The plates were then incubated with a modified RIPA buffer for 3 minutes and washed twice with PBS. Adhesion complexes were isolated using an adhesion recovery solution. To precipitate the adhesion complex proteins, four volumes of acetone were added, and the mixture was stored overnight at −80°C. The precipitated proteins were collected by centrifugation at 16,000 x g for 20 minutes at 4°C. The pellet was washed with acetone, dried in a fume hood at room temperature for about 20 minutes, resuspended in SDS-PAGE sample buffer, and boiled before being subjected to western blotting.

### Proteomic analysis

LAM-EV and Normal-EV proteome analysis was carried out by Creative Proteomics and the total of 2289 proteins were identified. The fold-change cutoff was set when proteins with quantitative ratios above 2 or below 1/2 are deemed significant. Proteins of relative quantitation were divided into two categories. Quantitative ratio over 2 was considered up-regulation while quantitative ratio less than 1/2 was considered down-regulation. Intensive bioinformatics analyses were carried out to analyze those quantifiable proteins, including GO annotation, KEGG annotation, cluster analysis, volcano plot, and protein-protein interactions analysis.

### RNA sequencing analysis

RNA sequencing analyses were carried out by Quick Biology and Novogene. Quick Biology analyzed SCID mice-based studies. For this, the reads were first mapped to the latest UCSC transcript set using Bowtie2 version 2.1.0^86^ and the gene expression level was estimated using RSEM v1.2.15^87^. TMM (trimmed mean of M-values) was used to normalize the gene expression. Differentially expressed genes were identified using the edgeR program^88^. Genes showing altered expression with p < 0.05 and more than 1.5-fold changes were considered differentially expressed. Goseq^89^ was used to perform the GO enrichment analysis and Kobas^90^ was used to performed the pathway analysis.

Novogene analyzed 621-101 sphere-based studies. For this, sample quantification, integrity and purity was checked by using Agilent 5400 instrument. Messenger RNA was purified from total RNA using poly-T oligo-attached magnetic beads. After fragmentation, the first strand cDNA was synthesized using random hexamer primers, followed by the second strand cDNA synthesis using either dUTP for directional library or dTTP for non-directional library^91^. The library was checked with Qubit and real-time PCR for quantification and bioanalyzer for size distribution detection. Quantified libraries were pooled and sequenced on Illumina platforms, according to effective library concentration and data amount. Raw data (raw reads) of fastq format were firstly processed through in-house perl scripts. In this step, clean data (clean reads) were obtained by removing reads containing adapter, reads containing ploy-N and low-quality reads from raw data. At the same time, Q20, Q30 and GC content the clean data were calculated. All the downstream analyses were based on the clean data with high quality. Reference genome and gene model annotation files were downloaded from genome website directly. Index of the reference genome was built using Hisat2 v2.0.5 and paired-end clean 1 reads were aligned to the reference genome using Hisat2 v2.0.5. The Hisat2 was selected as the mapping tool for that Hisat2 can generate a database of splice junctions based on the gene model annotation file and thus a better mapping result than other non-splice mapping tools^92^. The featureCounts v1.5.0-p3 was used to count the reads numbers mapped to each gene^93^. Then, FPKM of each gene was calculated based on the length of the gene and reads count mapped to this gene. FPKM, expected number of Fragments Per Kilobase of transcript sequence per Millions base pairs sequenced, considers the effect of sequencing depth and gene length for the reads count at the same time, and is currently the most commonly used method for estimating gene. (For DESeq2^94^ with biological replicates) Differential expression analysis^95^ of two conditions/groups (two biological replicates per condition) was performed using the DESeq2Rpackage (1.20.0). DESeq2 provide statistical routines for determining differential expression in digital gene expression data using a model based on the negative binomial distribution. The resulting P-values were adjusted using the Benjamini and Hochberg’s approach for controlling the false discovery rate. Genes with an adjusted P-value <=0.05found by DESeq2 were assigned as differentially expressed. (For edgeR^88^ without biological replicates) Prior to differential gene expression analysis, for each sequenced library, the read counts were adjusted by edgeR program package through one scaling normalized factor. Differential expression analysis of two conditions was performed using the edgeR R package (3.22.5). The P values were adjusted using the Benjamini & Hochberg method. Corrected P-value of 0.05 and absolute foldchange of 2were set as the threshold for significantly differential expression. Gene Ontology^89^ (GO) enrichment analysis of differentially expressed genes was implemented by the cluster Profiler R package, in which gene length bias was corrected. GO terms with corrected P-value less than 0.05 were considered significantly enriched by differential expressed genes. KEGG is a database resource for understanding high-level functions and utilities of the biological system, such as the cell, the organism and the ecosystem, from molecular-level information, especially large-scale molecular datasets generated by genome sequencing and other high-through put experimental technologies (http://www.genome.jp/kegg/). The clusterProfiler R package was used to test the statistical enrichment of differential expression genes in KEGG^96^ pathways. The Reactome database brings together the various reactions and biological pathways of human model species. Reactome pathways with corrected P-value less than 0.05 were considered significantly enriched by differential expressed genes. The DO (Disease Ontology) database describes the function of human genes and diseases. DO pathways with corrected P-value less than 0.05were considered significantly enriched by differential expressed genes. The DisGeNET database integrates human disease-related genes. DisGeNET pathways with corrected P-value less than 0.05 were considered significantly enriched by differential expressed genes. The clusterProfiler software was used to test the statistical enrichment of differentially expressed genes in the Reactome pathway, the DO pathway, and the DisGeNET pathway. Gene Set Enrichment Analysis (GSEA) is a computational approach to determine if a pre-defined Gene Set can show a significant consistent difference between two biological states. The genes were ranked according to the degree of differential expression in the two samples, and then the predefined Gene Set were tested to see if they were enriched at the top or bottom of the list. Gene set enrichment analysis can include subtle expression changes. The local version of the GSEA analysis tool was used (http://www.broadinstitute.org/gsea/index.jsp), GO, KEGG, Reactome, DO and DisGeNET data sets were used for GSEA independently.

### Rat DNA quantification

Rat DNA in SCID mice lungs was quantified as previously described^97^.

### FAP ELISA

Mouse plasma FAP levels were determined by solid phase sandwich ELISA according to the manufacturer’s instructions using DuoSet Mouse FAP (R & D Systems, USA #DY8647-05).

### Study approval

Human plasma samples from LAM patients and healthy donors were from the Center for LAM Research at Brigham and Women’s Hospital with obtained informed consent from all human participants under Institutional Review Board approval.

The following mouse strains were used: B6.Cg-Tg(Nes-cre)1Kln/J, Tsc1tm1Djk/J (from Jaxson Laboratory), SCID, NOD-Prkdc^em26Cd52^Il2rg^em26Cd22^/NjuCrl and CB17/Icr-*Prkdc^scid^*/IcrlcoCrl (from Charles River). Mouse studies were performed in compliance with the U.S. Department of Health and Human Services Guide for the Care and Use of Laboratory Animals and approved by the TTUHSC Institutional Animal Care and Use Committee (10034/10036/22006).

### Animal studies

For ELT3-based studies, female CB17/Icr-*Prkdc^scid^*/IcrlcoCrl (CB17 SCID/Fox Chase SCID) mice at 4-6 weeks of age were purchased from Charles River Laboratories. For 621-101-based studies, female NOD-Prkdc^em26Cd52^Il2rg^em26Cd22^/NjuCrl (NCG) mice at 7-8 weeks of age were purchased from Charles River Laboratories.

#### For short-term lung colonization using SCID mice and NT derived EV

The neural tube (NT) from an E15.5 embryo was collected from both *Nestin-Cre^+^Tsc1^−/−^* and wild-type littermates and cultured as described^9,74^. EV were purified from the culture media using UC method^9^, labeled with AF488 ExoGlow, and tail vein injected to SCID mice. 48 hours post EV inoculation 5 × 10^5^ of ELT3 cells were tail vein injected and mice were harvested 72 hours later.

#### For short-term lung colonization using NCG mice and plasma derived EV

5 × 10^5^ of 621-L9 and TSC2-add back cells in 100 µL of PBS were tail vein injected into NCG mice. Six hours later, mice were harvested and plasma EV were isolated using 30% sucrose cushion and UC method, and labeled with AF488 ExoGlow. Approximately equal amount of EV or EDP protein were tail vein injected to tumor-free NCG mice in the first experiment, whereas equal amount of EV protein and higher amount of EDP protein were tail vein injected in the second experiment. 72 hours post EV or EDP inoculation, mice were tail vein injected with 5 × 10^5^ of 621-L9 luciferase cells and imaged using IVIS bioluminescence imaging system as previously described^98^.

#### For short-term lung colonization using NCG mice and conditioning media derived EV

NCG mice were tail vein injected with 3 µg of EV, EDM, or PBS. 48 hours post EV, EDM, or PBS inoculation later mice were tail vein injected with 5 × 10^5^ 621-L9 luciferase cells and imaged using IVIS bioluminescence imaging system as previously described^98^.

#### In vivo bioluminescent reporter imaging

Ten minutes before imaging, mice were given D-luciferin (120 mg/kg, i.p., PerkinElmer Inc, 122799). Bioluminescent signals were recorded using the IVIS Spectrum System. Total photon flux of chest regions was analyzed and quantified.

#### In vivo MMP study

The IVISense MMP-750 FAST Fluorescent Probe (MMPSense) from Revvity was used following the manufacturer’s instructions. The probe was reconstituted in 1.2 ml PBS and tail vein injected into NCG mice 24 hours post EV inoculation at the dose of 2 nmol (100 µl) per mouse. The fluorescent signal was recorded using IVIS imaging system.

### Statistics

Data are expressed as mean ± SEM. Intergroup variations were assessed using either a two-tailed Student’s t-test, one-way ANOVA, or two-way ANOVA, as appropriate. Post-hoc comparisons were conducted with Tukey’s multiple comparison test. Statistical analyses were performed using GraphPad Prism, version 10.2.3. Differences were considered statistically significant at p-values < 0.05.

## Results

### EV biogenesis is increased and plasma EV cargo modified in LAM patients compared to healthy donors

To determine whether LAM patient-derived EV (LAM-EV) are different from EV from healthy donors (Normal-EV), we isolated EV from plasma of LAM patients and healthy donors, using ultracentrifugation and 30% sucrose method, and analyzed by direct light scattering (DLS), fluorescent activated cell sorting (FACS), and Western immunoblotting. LAM-EV were more frequent within size range of 0-50 nm compared to Normal-EV (Suppl. Fig. 1). EV fractions from both cohorts were negative for contaminants, including mitochondria (TFAM), endoplasmic reticulum (ER, GRP94, Calnexin), or apoptotic bodies (Annexin V) (Fig. 1A), and expressed EV markers CD63, CD81, and CD9 (Fig. 1A and 1B, Suppl. Fig. 1B). The endocytic origin of EV is supported by expression of Rab27A/B, flotillin2, and ALIX (Fig. 1A). Importantly, LAM-EV have increased expression of majority of EV-associated proteins, including Rab27A/B, ALIX, and CD9 (Fig. 1A and 1B), supporting increased EV biogenesis in LAM compared to healthy individuals.

**Figure 1.**
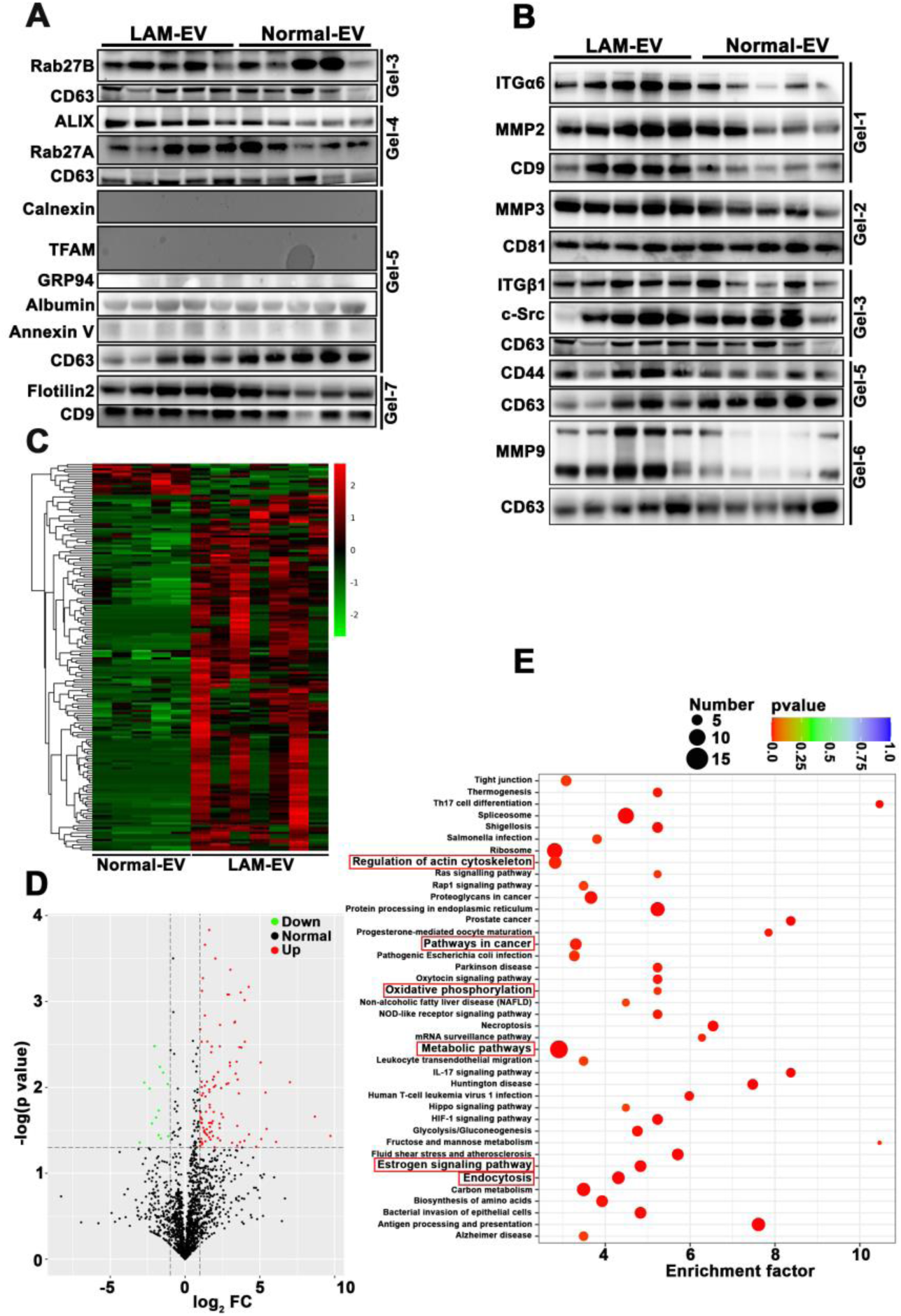
LAM-EV have modified cargo compared to Normal-EV. (A-B) Immunoblots of EV from plasma of LAM patients (LAM-EV) and healthy donors (Normal-EV), (n=5). (A) EV associated proteins and *JEV* controls; (B) Targeted analysis of EV cargo. (C-E) Proteome of LAM-EV (n=8) and Normal-EV (n=5; for 3 patients EV were isolated from different aliquots and analyzed separately). (C) The hierarchical clustering heat map of differentially expressed proteins in EVs; (D) Volcano plot; and (E) top 40 enriched pathways; The same EV loading controls were used in panel A and B as data were split in between these panels.

Since pulmonary LAM is a low-grade metastasizing sarcoma^1^ and EV-derived integrins regulate tumor cells organotropism^33,35^ with lung-tropic EV-ITGα6/β1/β4^35^ binding to the lung-resident fibroblasts and epithelial cells to promote lung metastasis^35^, we examined the expression of ITGα6/β1 in LAM-EV. We also assess LAM-EV expression of several metalloproteinases, CD44, and c-Src, as all are implemented in LAM pathogenesis^64,99–103^. LAM-EV have increased expression of ITGα6/β1, MMP2, MMP3, MMP9, c-Src, and CD44 compared to Normal-EV (Fig. 1B), suggesting possible role of LAM-EV in lung tropism and disease progression. To gain insights into functions of LAM-EV, we compared the proteome of LAM- and Normal-EV. The total of 2289 EV proteins were identified, 149 and 13 were upregulated or downregulated in LAM-EV relative to Normal-EV, respectively (Fig. 1C and1D). Kyoto Encyclopedia of Genes and Genomes (KEGG) pathway analysis identified top 40 enriched pathways for differentially expressed proteins (DEP) in LAM-EV, including regulation of actin cytoskeleton, pathways in cancer, oxidative phosphorylation, metabolic pathways, estrogen signaling pathway, and endocytosis (Fig. 1E).

### The loss of TSC1/2 alters biochemical and physical characteristics of EV

To corroborate clinical data, we determined the impact of TSC1/2 loss on biochemical and physical EV properties and on EV cargo. EV from 621-101 (TSC-null EV) and from TSC2 addback cells (TSC2 EV)^9^ (Suppl. Fig. 2A) were isolated and characterized. The particle concentration of TSC-null EV and TSC2 EV, isolated by ultracentrifugation (UC) alone and analyzed by NTA, was 3.2×10^9^ and 2.9×10^9^ (particles/ml), respectively (Suppl. Fig. 2B-i). The mean size of TSC-null EV and TSC2 EV was 121.1 ± 1.8 nm and 142 ± 1.3 nm, respectively. The size distribution analysis indicated that TSC2 EV were more frequent within size range of 150-299 nm compared to TSC-null EV (Suppl. Fig. 2B-ii). The size distribution analysis by DLS of EV isolated by ultrafiltration (UF)^76^ and size exclusion chromatography (SEC) indicated that TSC-null EV were more frequent within size range of 100-150 nm compared to TSC2 EV (Suppl. Fig. 2C-i). Thus, loss of TSC2 alters particle concentrations, size, and size distribution of EV with TSC2-null EV being more concentrated and smaller than EV from TSC2 addback cells. Zeta potential analysis indicated negative charge of EV confirming the lack of aggregates and preservation of functionality (Suppl. Fig. 2C-ii). Both EVs express CD63 and CD9 according to FACS analysis (Suppl. Fig. 2D). The transmission electron microscopy (TEM) revealed cup-shaped morphology of these EV (Suppl. Fig. 2E). Western immunoblotting of EV and *JEV* controls^72^, loaded in the equal protein quantities, confirmed EV expression of tetraspanins, CD63, CD9, and CD81 (Fig. 2A). These EV were negative for contaminants such as albumin, mitochondria (TFAM), endoplasmic reticulum (ER, GRP94, Calnexin), or apoptotic bodies (Annexin V) (Fig. 2A). The endocytic origin of EV was supported by expression of Rab27A/B, ALIX, and flotillin-1/2 (Fig. 2A). Similar to LAM EV, TSC-null EV have increased expression of majority of EV-associated proteins, including ALIX, Rab27B, CD63, CD81, flotillin1/2, and CD9 (Fig. 2A), indicating that loss of TSC2 increases EV biogenesis.

**Figure 2.**
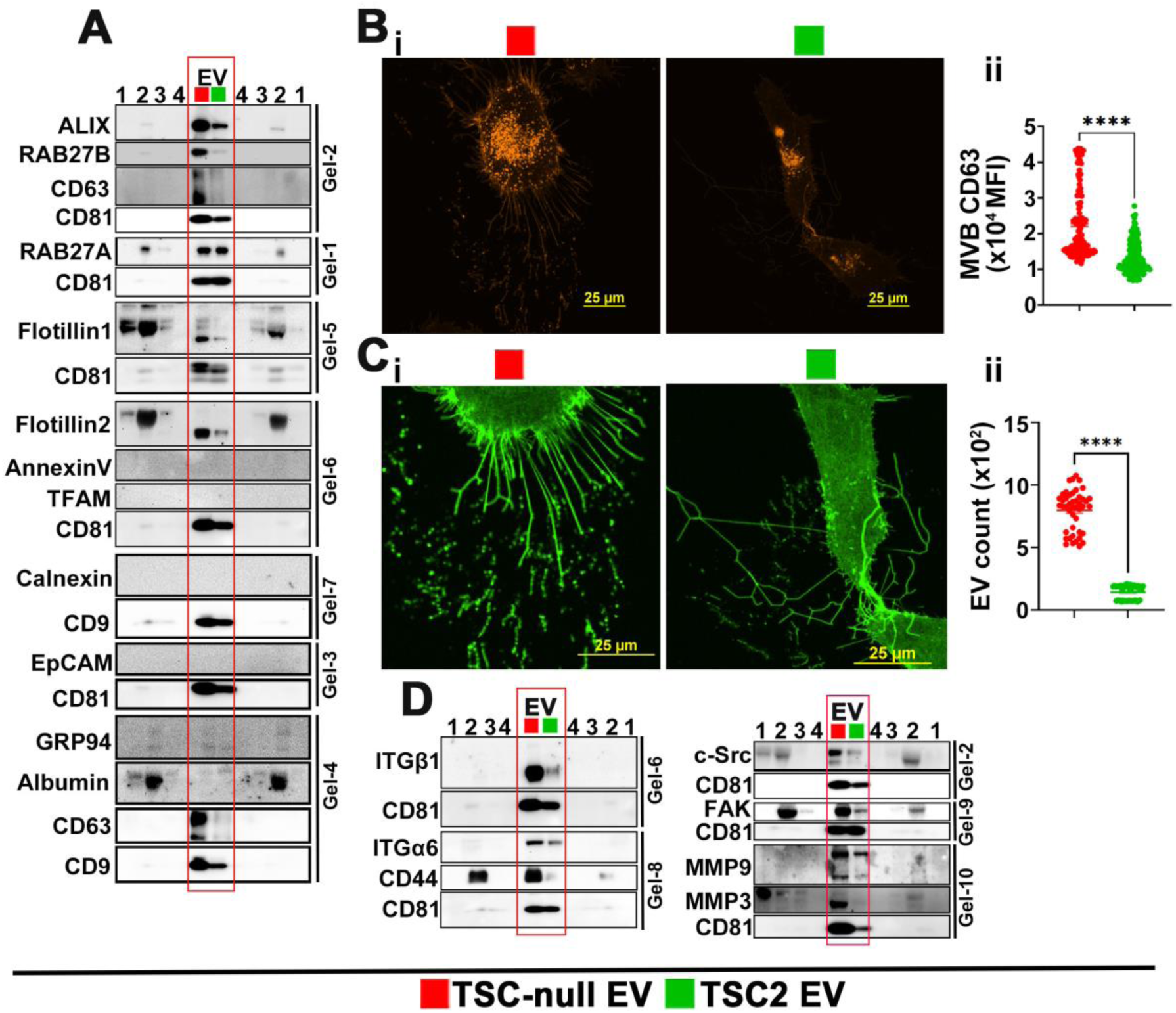
Loss of TSC1/2 increase EV biogenesis. (A) Representative immunoblots of EV associated proteins and *JEV* controls from conditioning media of 621-101 and TSC2 addback cells (n=2). (B-C) Immunofluorescence of adherent 621-101 cells expressing pHluo_M153-CD63-mScarlet; (B-i) red fluorescence indicates CD63 MVB expression, (B-ii) quantification of B-i; (C-i) green fluorescence indicates CD63**+** EV deposited on ECM, (C-ii) quantification of C-i. Data are Mean ± SEM of 3 independent experiment. (D) Representative immunoblots of targeted EV protein expression and *JEV* controls from conditioning media of 621-101 and TSC2 addback cells (n=3). ****P<0.0001by t-test. The same EV loading controls were used in panel A and D as data were split in between these panels.

To track CD63^+^ EV, we used CD63 dual-color reporter pHluo_M153-CD63-mScarlet^75^ in 621-101 and TSC2 addback cells. This construct exhibits red fluorescence under acidic (e.i. in multivesicular bodies (MVB) or dual (green and red) fluorescence in neutral conditions (e.i. in secreted EV)^75^. TSC-null cells have increased intracellular/MVB expression of CD63 compared to TSC2 addback cells (Fig. 2B), suggesting increased CD63 sorting to TSC-null MVB and EV. Since TSC2 loss affects biogenesis of fluid phase EV (Fig. 2A and Suppl. Fig. 2), we examined impact of TSC2 loss on EV deposited on extracellular matrix (ECM) using the same reporter. The loss of TSC2 increases EV deposition on ECM compared to TSC2 addback cells (Fig 2C).

### TSC-null EVs are enriched with ITGα6/β1, c-Src, and FAK

Since EV-ITGα6/β1 promote lung metastasis^35^ and metalloproteinases, CD44, and c-Src are implicated in LAM pathogenesis^64,99–103^, we examined the expression of these proteins in TSC-null and TSC2 EV. EV were isolated from adherent 621-101 cells, cultured for 72 hours, using UF^76^ and SEC. Similar to LAM EV, TSC-null EV are enriched with ITGα6/β1, CD44, c-Src, FAK, MMP9, and MMP3 compared to TSC2 EV (Fig. 2D). Overall our data suggest that TSC1/2 regulates EV cargo sorting of soluble and membrane EV proteins.

### TSC-null EV enhance migration and invasion of 621-101 adherent cells

LAM is a metastatic sarcoma^1–4^ and cell migration and stromal invasion are essential for metastatic progression, therefore, we determined the impact of EV isolated from adherent 621-101 and TSC2 addback cells, incubated for 72 hours, on 621-101 adherent cells’ migration and invasion, using scratch and transwell assays. TSC-null EV increase 621-101 cells’ migration to a greater extent than TSC2-2-EV (Fig. 3A and Suppl. Fig 3A), which is associated with increased expression of ITGα6/β1, activation of c-Src, indicated by increased Y416 phosphorylation, c-Src- and integrin-mediated activation of FAK, indicated by increased phosphorylation of Y576/577 and Y397, respectively (Fig. 3B). The c-Src-FAK axis increased activation is associated with an increased activation of AKT, indicated by S473 phosphorylation (Fig. 3B). In addition, we found increased actin polymerization and paxillin activation, as indicated by increased F-actin expression and Y118 phosphorylation, respectively (Fig. 3C), as well as increased co-localization of both proteins in cell treated with TSC-null EV (Fig 3C). TSC-null EV also increased invasion of 621-101 cells compared to TSC2-EV (Fig. 3D). The treatment of 621-101 cells with inhibitors of EV uptake or biogenesis (Suppl. Fig. 3B-D) prevents TSC-null-EV mediated increase in 621-101 cells migration (Fig. 3E and Suppl. Fig. 3F, G) and invasion (Fig. 3F and Suppl. Fig. 3E, H). Similarly, the inhibition of c-Src in these cells (Suppl. Fig. 3I) prevents TSC-null-EV mediated increase in 621-101 cells migration (Fig. 3G) and invasion (Fig. 3H). Thus, these data indicate that TSC-null EV increase 621-101 cell migration and invasion via the EV-ITGα6/β1-c-Src-FAK-AKT regulatory axis.

**Figure 3.**
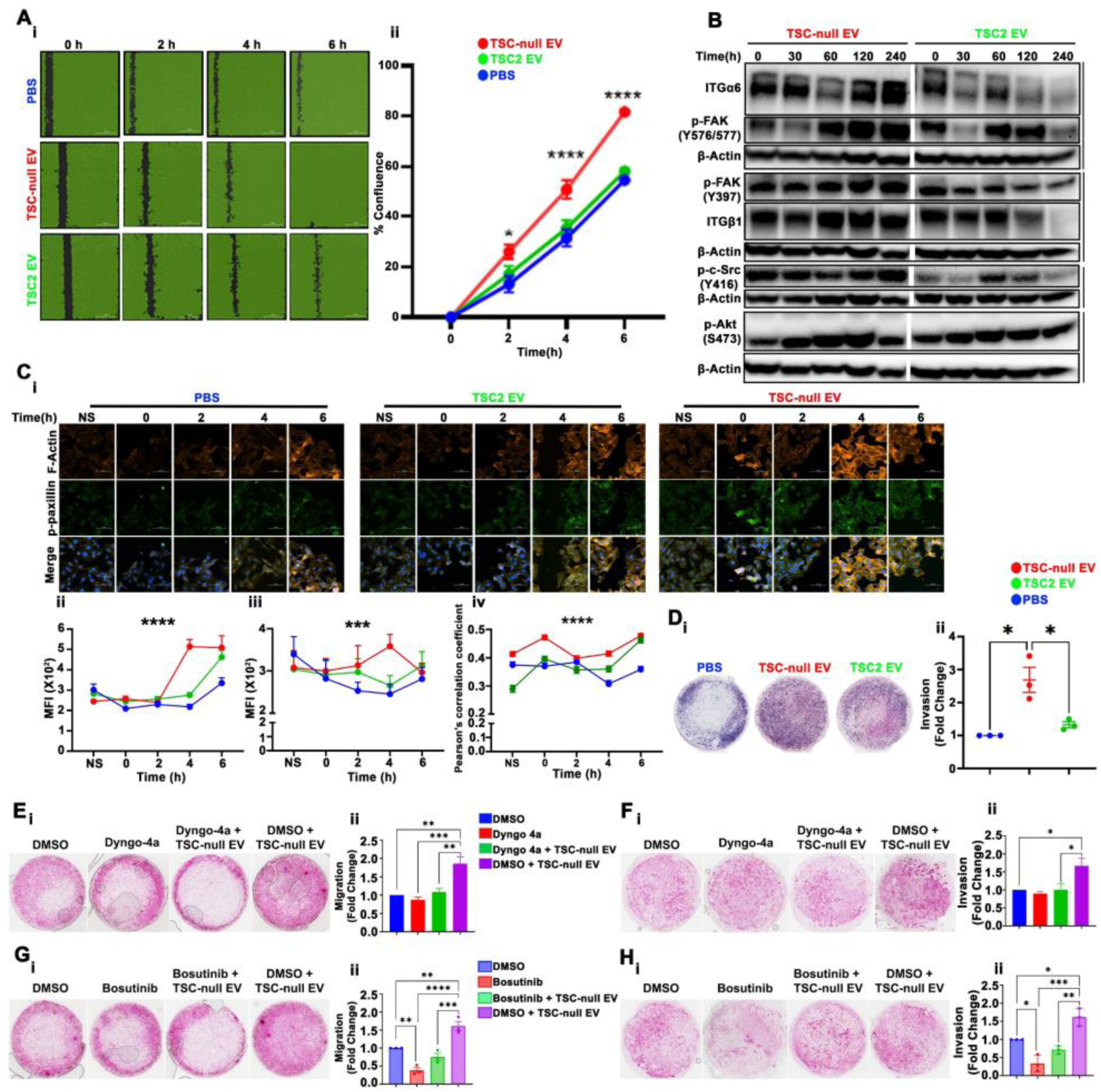
TSC-null EV enhance migration and invasion of adherent TSC-null cells. (A) 621-101 cells were seeded and treated with EV and when approximately 95% confluent scratched; (A-i) Representative mark-up images of migrated cells. (A-ii) Quantification of wound closure at indicated timepoint with respect to the wound area at time 0. (B) Expression of ITG-α6/β1, phospho-FAK, phospho-c-Src, and phospho-Akt in migrated 621-101 cells treated with TSC-null EV *vs.* TSC2 EV by immunoblot. β-actin was used as a loading control. (C) Immunofluorescence of (C-i) F-Actin and p-paxillin in migrated 621-101 cells treated with TSC-null EV *vs.* TSC2 EV, (C-ii), quantification of F-Actin and (C-iii) p-paxillin, (C-iv) co-localization of F-Actin and p-paxillin. (D) Transwell invasion assay of adherent 621-101 cells treated with TSC-null EV *vs.* TSC2 EV; (D-i) Representative images and (D-ii) quantification of D-i as a fold change in number of cells relative to PBS. (E-H) adherent 621-101 cells treated with vehicle or inhibitor of EV uptake (E-F), or vehicle or inhibitor of c-Src (G-H) in the presence or absence of TSC-null EV; (E, G) Transwell migration assay; (E-i, G-i) Representative images and (E-ii, G-ii) quantification of E-i and G-i as a fold change in number of cells relative to DMSO (E-ii, G-ii). (F, H) Transwell invasion assay (F-i, H-i) Representative images and (F-ii, H-ii) quantification of F-i, and H-i as a fold change in number of cells relative to DMSO. Data are representative of 3 independent experiments. Graphs plotted show Mean ± SEM from 3 independent experiments. *P<0.01; **P<0.001; ***P<0.0001; ****P<0.0001by one-way ANOVA with Tukey’s multiple comparison test.

### TSC-null EV enhance CSCs and metastable phenotypes of 621-101 spheres

TSC-null EV and TSC2 EV were isolated from adherent 621-101 or TSC2 addback cells, respectively, grown for 3 or 7 days representing nutrient rich- or nutrient low-like environment, respectively. Under these experimental conditions, these EV mimic EV released from primary tumor (primary tumor like EV) cells. To generate TSC-null EV and TSC2 EV mimicking EV released from metastasizing LAM cells (metastasizing tumor-like EV), we isolated EV from 621-101 or TSC2 addback spheres, respectively, grown for 7 days in ultra-low attachment plates. Impact of these EV subtypes on LAM (621-101) cell CSC-like phenotypes was determined using primary and secondary sphere formation, proliferation, aldehyde dehydrogenase activity (ALDH), and sphere cell migration and invasion assays. TSC-null EV subtypes increase CSCs properties of 621-101 spheres to a greater extent than TSC2 EV, as indicated by increased diameter of primary spheres (Fig. 4A), ability to form secondary spheres (Fig. 4B), cell proliferation (Fig.4C), ALDH activity, (Fig. 4D), increased sphere cell migration (Fig. 4E), and invasion (Fig.4F). Interestingly, metastasizing-like TSC-null EV led the greater increase in sphere size, ALDH activity, and sphere cell migration compared to primary-like tumor TSC-null EV subtype (Fig. 4A and 4D-E). The treatment of 621-101 spheres with inhibitors of EV uptake or biogenesis (Suppl. Fig. 4A) resulted in smaller (Fig. 4G, 4H, and Suppl. Fig. 4B-ii) and less migrating spheres (Suppl. Fig. 4C), respectively. Similar to inhibitors of EV uptake and biogenesis, the inhibition of c-Src in these spheres (Suppl. Fig. 4E) resulted in smaller (Fig. 4I) and less migrating spheres (Suppl. Fig. 4D). Collectively, these data suggest different functions for different TSC-null EV subtypes with sphere-derived EV having the greatest potential to enhance CSCs and metastable phenotypes of LAM cells.

**Figure 4.**
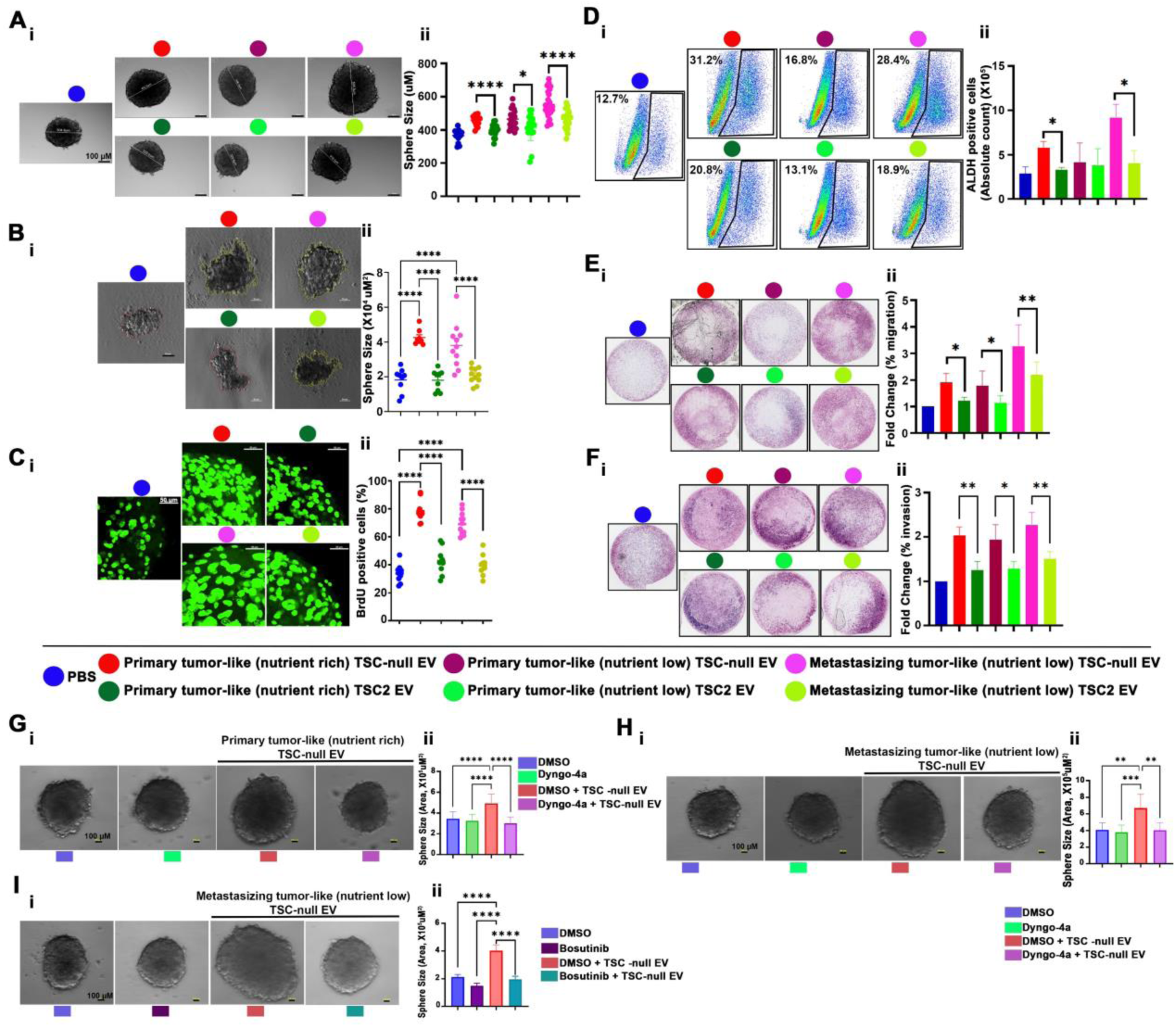
TSC-null EV enhances CSCs and metastable phenotypes of 621-101 spheres. (A-B) Sphere assay; (A) Primary sphere and (B) secondary sphere assay in 621-101 cells incubated with indicated EV subtypes; Representative images (A-i, B-i) and (A-ii, B-ii) quantification of sphere size. (C) Immunofluorescence of BrdU-positive 621-101 cells incubated with indicated EV subtypes. The image is from single Z stack of 1 µm each. (D) ALDEFLUOR assay; (D-i) FACS analysis of ALDH-positive 621-101 sphere cells incubated with indicated EV subtypes, (D-ii) The absolute count of ALDH-positive cells is represented in the bar graph (n=4 for PBS group, n=3 for all remaining). (E-F) Transwell migration and invasion assay of 621-101 spheres incubated with indicated EV subtypes; (E-i, F-i) Representative images, and (E-i, F-ii) quantification of E-i and F-i as a fold change in number of cells relative to PBS. (G-I) Sphere assay in 621-101 cells incubated or not with indicated EV subtypes in the presence of (G-H) DMSO or EV uptake inhibitor, Dyngo4a (10 µM), or (I) c-Src inhibitor, Bosutinib (1 µM). (G-I-i) Representative images and (G-I-ii) quantification of sphere sizes. Data are represented as Mean ± SEM (n=3). Statistical significance was determined using t-test (D) and one-way ANOVA test. *p < 0.05, **p < 0.01, ***p < 0.001, ****p < 0.0001.

### Shuttling ATP synthesis to pseudopodia and activation of integrin adhesion complex signaling drive TSC-null EV subtypes mediated CSC metastable phenotypes of LAM cells

To understand the different mode of action of different TSC-null EV subtypes, RNA-Seq of 621-101 spheres treated with primary-like or metastasizing tumor-like EV was performed. 621-101 spheres treated with primary tumor-like TSC-null EV upregulated and downregulated 805 and 297 genes, respectively, relative to spheres treated with TSC2 EV (Fig. 5A and 5B). Top upregulated genes are involved in the regulation of oxidative phosphorylation and ROS (Fig. 5B). RT-qPCR confirmed upregulation of OXPHOS related genes, including *NDUFB4*, *NDUFB5*, *COQ3*, *MRLP22*, and *MRLP48* (Fig. 5C) and this upregulation is associated with increased levels of ATP (Fig. 5D). The upregulation of OXPHOS genes and increase in ATP were likely mediated by increased expression and delivery of critical mitochondrial function regulator, Nrf2^104^ by primary tumor -like TSC-null EV, compared to TSC2 EV (Fig. 5E). Clinical significance of these data is underscored by Nrf2 enrichment in patient LAM-EV compared to Normal-EV (Fig. 5F). In addition, we found increased expression of Nrf2, p-AMPK, ITGβ1, MMP14, MMP2, p-FAK, and p-AKT in TSC-null EV vs TSC2 EV treated spheres (Fig 5G). Cumulatively, these data suggest that primary tumor-like TSC-null EV shift metabolism of sphere cells toward OXPHOS, to enhance sphere cell migration, and are consistent with AMPK function in mitochondria trafficking to the leading edge and protrusive structures of the cell during migration and invasion^84^. Indeed, the assessment of subcellular energetics by measuring ATP in chemotactic (FBS) pseudopodia (Pd) and cell bodies (CB), using transwell-like cell culture inserts^84^, revealed higher levels of ATP, p-AMPK, ATP synthase, TFAM, activated FAK, and c-Src in Pd compared to CB of spheres treated with primary tumor -like TSC-null EV (Fig. 5H and5I), suggesting EV mediated shuttling of ATP synthesis to Pd. Analysis of Pd and CB of spheres treated with TSC2 EV showed reversed phenotypes (Fig 5I). The TSC-null EV mediated increase in Pd localized ATP synthesis upregulates ITGβ1, CD44, and MMP9, and activates paxillin, FAK, and c-Src indicated by their phosphorylation in migrated sphere cells (Fig. 5J). Furthermore, real-time cell tracking approach confirmed TSC-null EV-mediated increase in accumulated distance and velocity of migrated sphere cells (Fig. 5K). These data suggest that primary tumor-like TSC-null EV mediate Pd localized ATP synthesis to promote sphere cell migration via activation of the ITG-c-Src-FAK axis.

**Figure 5.**
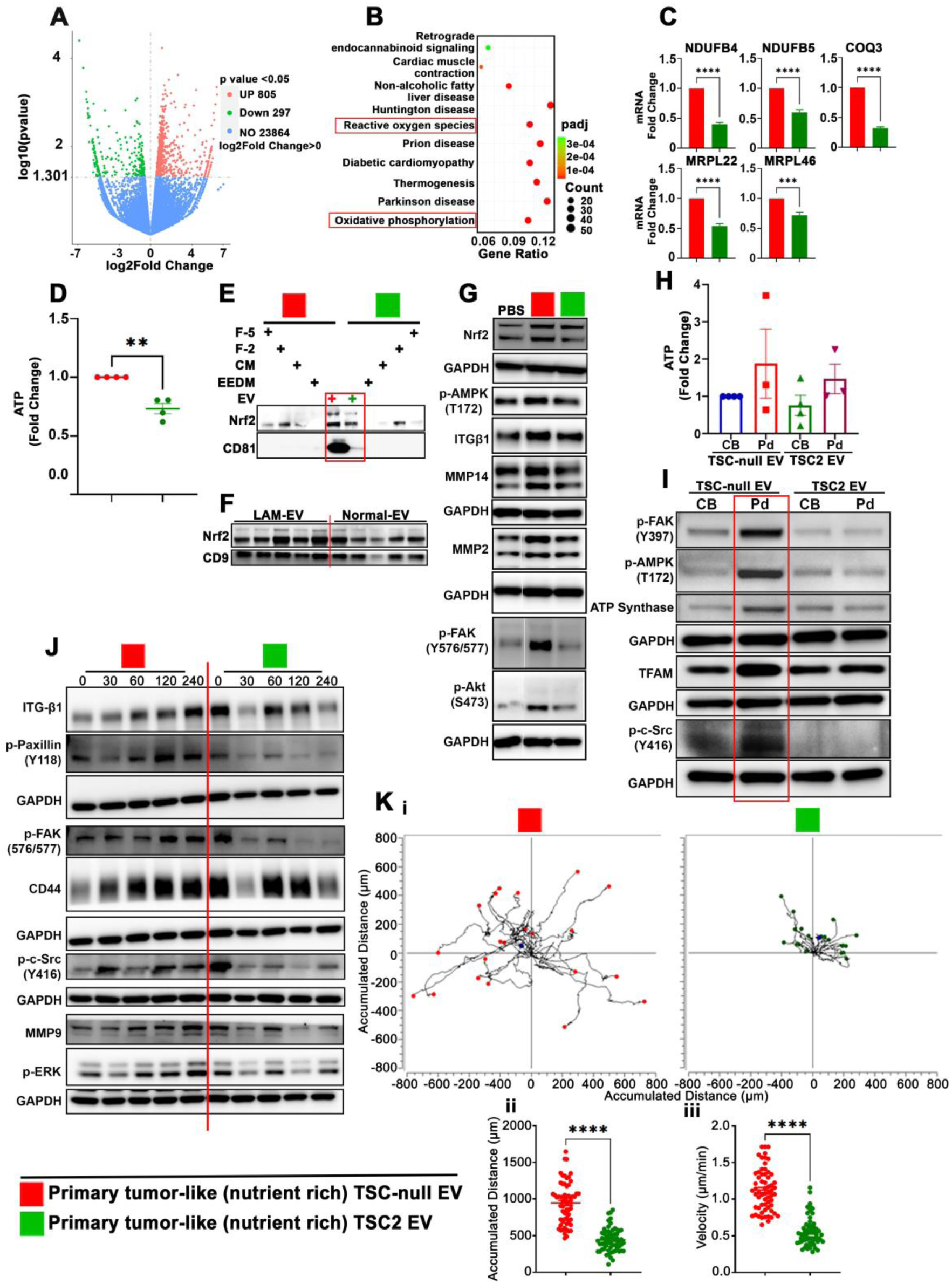
Primary tumor-like TSC-null EV promotes pseudopodia localized ATP synthesis to increase TSC-null sphere cell migration. (A) Volcano plot of differentially expressed genes in 621-101 spheres treated with primary-like TSC-null EV in comparison to TSC2 EV (n=3). (B) Top enriched pathways from (A). (C) Validation of RNA-seq data for Oxidative phosphorylation genes by RT-qPCR (n=3). (D) Cellular ATP level in 621-101 spheres treated with indicated EV (n=4). (E-F) Immunoblot of TSC-null EV *vs* TSC2 EV and LAM-EV *vs* Normal EV, respectively. The loading control from panel F was also used in Figure 1A-B. (G) Immunoblot of 621-101 spheres treated with PBS or primary tumor-like EV. (H) Relative levels of ATP (per microgram of protein) were assayed from equal amounts of extracts from purified CB and Pd (n=3-4). (I) The protein levels of phospho-T172 AMPK, phospho-Y397 FAK, phospho-Y416 Src and ATP synthase in CB and Pd of 621-101 sphere cells treated with indicated EV were assessed by immunoblot (n=3). (J) Immunoblot of migrated 621-101 spheres cells treated with primary tumor-like TSC-null *vs* TSC2 EV. (K) Time-lapse imaging of migrated 621-101 sphere cells. (K-i) Trajectory plots, (K-ii) accumulated distance and (K-iii) the moving velocity; (n=60 cells from 3 independent experiment). Data are representative of 3-4 independent experiments. Bars show mean ± SEM. **P<0.001; ***P<0.0001; ****P<0.0001by unpaired t-test.

Metastasizing tumor-like TSC-null EV upregulated and downregulated 100 and 99 genes in 621-101 spheres, respectively, relative to spheres treated with TSC2 EV (Fig. 6A and 6B). Top upregulated genes are involved in ECM receptor interaction (Fig. 6B). RT-qPCR analyses confirmed upregulation of ECM related genes, including *TNC*, *MMP3*, *FN1*, *HIST1H3I*, and *ABI3BP* (Fig 6C). The increase in ECM gene expression in TSC2-null EV treated spheres compared to TSC2 EV treated spheres associates with moderate increase in the expression of ITGα6/β1, MMP3, CD44, as well as increased activation of c-Src, FAK, ERK, and AKT indicated by their phosphorylation (Fig. 6D). TSC-null EV mediated whole-cell increase in expression of ITGα6/β1, CD44, talin, paxillin, ILK, and vinculin, as well as c-Src, and FAK activation suggests that these EV regulate the formation of integrin adhesion complexes (IAC)^105,106^. Consistently, we found increased formation of IAC in migrating sphere cells treated with TSC-null EV (isolated as in reference^106^), demonstrated by increased expression of ITGα6/β1 and canonical IAC proteins^105^ including talin, vinculin, paxillin, ILK, c-Src, FAK, and tetraspannins CD63 and CD9 (Fig. 6E). The TSC-null EV mediate increase in IAC formation upregulates vinculin and ITGα6/β1, and activates paxillin, FAK, and c-Src, indicated by their phosphorylation in migrated spheres (Fig. 6F). The TSC-null EV mediated upregulation of IAC signaling was likely interceded by increased expression of vinculin, paxillin, and ILK in metastasizing tumors-like TSC-null EV, compared to TSC2 EV (Fig. 6G). These data were corroborated by paxillin and ILK enrichment in LAM-EV compared to Normal-EV (Fig. 6H). Furthermore, real-time cell tracking approach confirmed TSC-null EV mediated increase in accumulated distance and velocity of migrated sphere cells (Fig. 6I). These results suggest that metastasizing tumor-like TSC-null EV promote sphere cell migration via increase in IAC formation, which is mediated by the delivery of pre-formed building blocks of ILK, vinculin and paxillin heterodimers, and IAC signaling.

**Figure 6.**
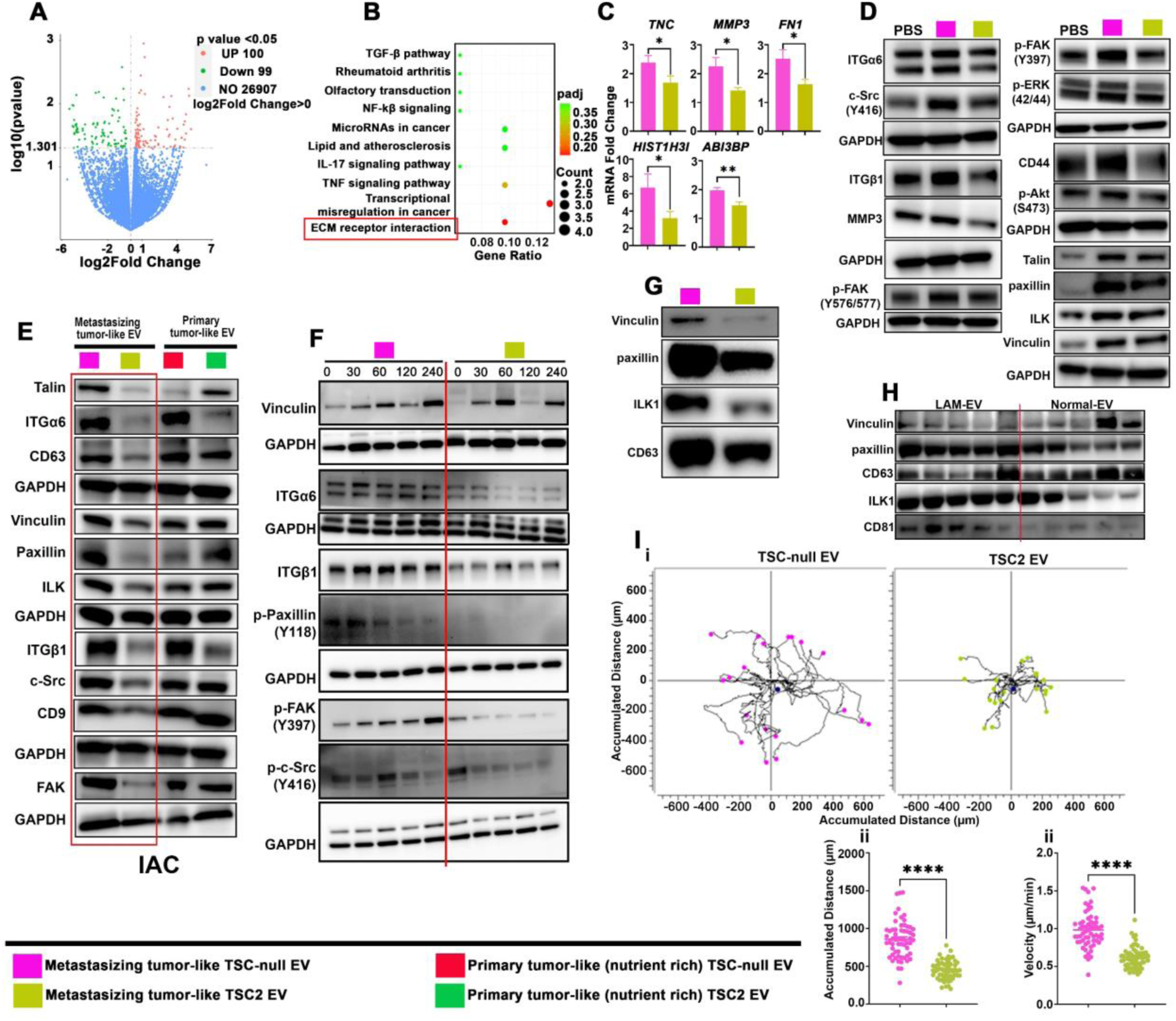
Metastasizing tumor-like TSC-null EV promote sphere cell migration via increase in IAC formation. (A) Volcano plot of differentially expressed genes in 621-101 spheres cells treated with metastasizing tumor-like TSC-null EV in comparison to TSC2 EV (n=3). (B) Top enriched pathways from (A). (C) Validation of RNA-seq data for ECM receptor genes by RT-qPCR (n=3). (D) Immunoblot of 621-101 spheres treated with PBS or metastasizing tumor-like EVs. (E-F) Immunoblot of IAC isolated from 621-101 spheres treated with indicated EV (E), and migrated sphere cells (F). (G-H) IAC protein expression in (G) metastasizing tumor-like TSC-null EV compared to TSC2 EV, or (H) LAM-EV and Normal-EV by immunoblot. (I) Time-lapse imaging of migrated 621-101 sphere cells treated with indicated metastasizing tumor-like EV; (I-i) Trajectory plots, (I-ii) accumulated distance and (I-iii) the moving velocity; (n=60 cells from 3 independent experiment). Data are representative of 3 independent experiments. Bars show mean ± SEM. *P<0.01; **P<0.001; ****P<0.0001by unpaired t-test.

### EV from TSC-null cells increase lung metastatic burden in a mouse model of LAM

CD9^+^CD81^+^CD63^+^EV from *Tsc1*-null or EV from wild type E15.5 mouse embryo neuronal progenitors^9^ (Suppl. Fig. 5A, 5C and 5D) were labeled and injected into the tail vein of female SCID mice 48 hr. prior to the i.v. injection of 0.5×10^6^ rat ELT3 cells (a well-characterized mouse model of LAM^98,107^). The 72 hr. after ELT3 cell injection, we found more rat DNA, reflecting metastatic burden, in the lungs of *Tsc1-*null vs. wild type EV injected mice (Fig. 7A and Suppl. Fig. 5B). The RNA-Seq/GO/Reactome/Kegg pathway analyses of these lungs revealed the upregulation and downregulation of 521 and 287 genes, respectively, in *Tsc-1* null EV-vs. wild type EV-injected mice (Figure 7B-C). *Tsc1*-null EV impacted genes involved in the regulation of ECM and collagen degradation, collagen biosynthesis, and modifying enzymes, as well as collagen fiber assembly (Fig. 7D). These data were corroborated by RT-qPCR and immunohistochemistry, demonstrating the increased expression of *Col1a1*, *Mmp14*, *Cxcl5*, and *Mmp2*, (Fig. 7E), collagen deposition (Fig.7F, blue color in histology images), and increased S100A4 in the lungs of *Tsc1*-null EV compared to wild type EV treated mice (Fig. 7G). Because EV-ITGβ1/α6 activates S100A4 in lung resident cells^35^, we examined expression of ITGβ1/α6 in *Tsc-1* null and wild type EV. *Tsc-1* null EV are enriched with ITGβ1/α6 relative to EV from wild type progenitors (Suppl. Fig. 5D), consistent with the roles of these ITGs in the activation of lung fibroblasts^35^ and S100A4 in lung resident cells^35^. Next, we isolated CD9^+^CD63^+^EV from SCID/NOD mice i.v. injected with LAM patient-derived 621L9 or TSC2 addback cells 6hr. prior to EV isolation (TSC-null EV *vs.* TSC2 EV) (Suppl. Fig. 5E and 5F). The treatment of tumor-free SCID/NOD mice with these plasma isolated TSC-null EV but not with TSC2 EV or EV-depleted plasma reduces the clearance of 621L9 cells, injected 72 hours post EV inoculation, from the lungs (Fig. 7H) and associates with increased expression of ECM, airway epithelial alveolar type 1/2, and fibroblast related genes, including *Itgβ1*, *Col11a*, *Mapk13*, *Cstk*, (ECM), *Abca3, Lrrc23* (epithelial) and *S100A4* (fibroblasts) in the lungs (Fig. 7I). The epithelial genes’ expression is consistent with gene enrichment in patient LAM-associated airway epithelial, alveolar type 1 and 2^4^. We also found increased fibroblast activating protein (FAP) in plasma of these mice (Fig. 7J).

**Figure 7.**
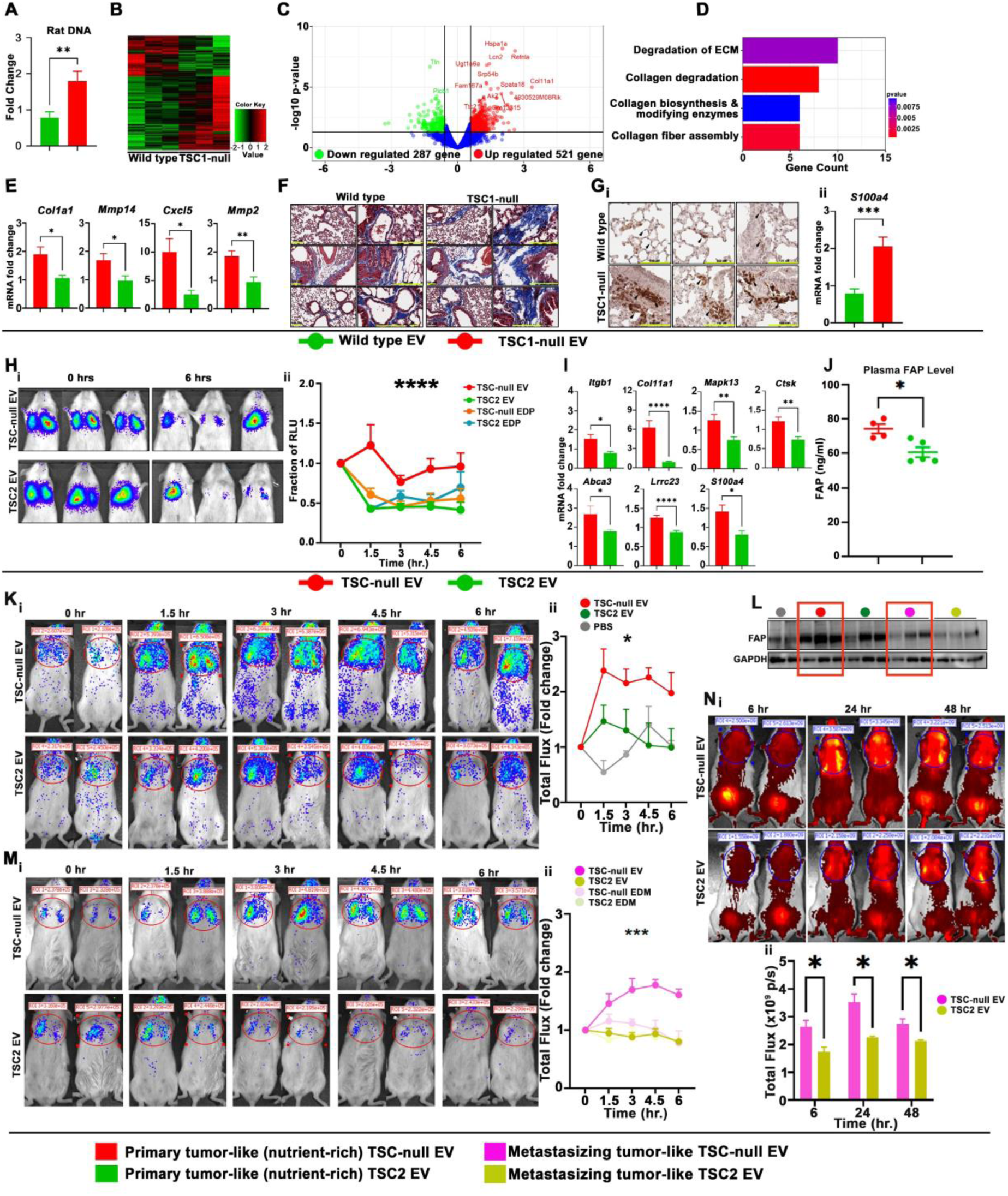
TSC-null EV promote lung seeding by LAM cells. (A) Rat DNA by qPCR. (B-D) RNA-Seq analyses: (B) heat map, (C)Volcano plot, and (D) top enriched pathways. (E) Expression of ECM genes by RT-qPCR. (F) Masson-Trichrome stain for collagen (blue). (G) Expression of S100A4 by (G-i) immunohistochemistry and (G-ii) RT-qPCR. (H) *In vivo* bioluminescent reporter imaging; (H-i) Bioluminescence imaging of the lungs from mice injected i.v. with 621-L9 cells 72 hr. after i.v. injection of 621L9- or TSC2-plasma EV, or controls injection and (H-ii) quantification of relative luciferase unit at different time points normalized to time zero (significance for “column factor”). (I) ECM, epithelial, and fibroblast related genes by RT-qPCR. (J) Plasma FAP ELISA. (K, M) Bioluminescence imaging of the lungs from mice injected i.v. with 621-L9 cells 72 hr. after (K-i) primary tumor-like TSC-null-EV or TSC2-EV, or (M-i) metastasizing tumor-like TSC-null-EV or TSC2-EV *vs* controls. (K-ii, M-ii) Quantification of relative luciferase unit at different time points normalized to time zero (significance for “interaction and column factor”). (L) The lung expression of FAP from mice treated with indicated EV subtypes. (N) Lung MMP activity using IVISSense MMP 750 FAST fluorescent probe from mice i.v. treated with metastasizing tumor-like EV 24 hours prior to probe injection (red boxes mark FAP expression in primary-*vs.* metastasizing tumor-like TSC-null EV); (N-i) Representative fluorescent images at 6, 24, and 48 hours post-MMP probe injection, and (N-ii) quantification of fluorescence photon flux in the chest region (n=4-5/group). *P < 0.05, **P < 0.01, ***P < 0.001. (B-ii) unpaired t test. (C-ii, D-ii) Two-way ANOVA with Tukey’s multiple comparison test.

The different functions of different TSC-null EV subtypes are supported by improved lung seeding by 621L9 cells (Fig. 7K) and greater expression of FAP (Fig. 7L) in the lungs of tumor-free SCID/NOD mice injected i.v. with primary tumor-like TSC-null EV compared to mice injected with metastasizing tumor-like TSC-null EV (Fig. 7M). In contrast, metastasizing tumor-like EV facilitated greater activation of MMPs compared to primary tumor-like EV (Fig. 7N, Suppl. Fig. 5G), indicating different and EV subtype dependent mechanism facilitating lung seeding by LAM cells. Although both TSC-null EV subtypes improved lung seeding by 621L9 cells compared to TSC2 EV, EDP, or PBS, (Fig. 7K and 7M), primary tumor-like EV are more efficient in supporting LAM cell retention in the lungs, suggesting greater contribution of this EV subtype to facilitating lung metastases. In summary, TSC-null EV subtypes derived from genotypically identical tumor cells, have different cargo-dependent functions that are linked to different mechanisms of cargo sorting. Engagement of these different mechanisms likely depends on differences in the tumor microenvironment (i.e. primary tumor cells/EV vs. metastasizing cells/EV).

## Discussion

This work provides evidence that LAM progression is, at least partially, mediated by increased biogenesis of LAM-EV and their enrichment with proteins known to drive lung organotropic metastases in breast cancer, including ITGα6/β1^35^. In addition, LAM-EV are enriched with several metalloproteinases, c-Src, and CD44, which is consistent with the dysregulation of these proteins in LAM and their roles in LAM progression^99–103^. KEGG enrichment analyses of LAM-EV identified top enriched pathways for DEP, including regulation of actin cytoskeleton, pathways in cancer, oxidative phosphorylation, metabolic pathways, estrogen signaling pathway, and endocytosis, supporting involvement of LAM-EV in LAM progression. The analysis of TSC-null EV and TSC addback EV demonstrated that loss of TSC2 increases EV biogenesis and alters physical and biochemical properties of EV. The loss of TSC2 mediates also changes in EV cargo sorting. Collectively, our data demonstrate that LAM-EV from patients share several features with TSC-null EV from LAM surrogate cells and that loss of TSC1/2 increases CD63+ LAM-EV biogenesis and modifies EV cargo. Our data are consistent with previous reports indicating that ITG-β1 is enriched in EV of melanoma cells^108^ and that loss of TSC1/2 increases EV biogenesis^9,10,109,110^, Despite the impact of Tsc1/2 deficiency on EV biology demonstrated by our data, the long-term rapamycin treatment had no conclusive effect on EV biogenesis^10^. In contrast, activation of mTOR inhibits EV release in *Tsc1/2*-null mouse embryonic fibroblasts and in hepatocytes^111^, which suggests that mTOR regulates EV release in different manner depending on cell type, physiological vs. pathological, and culture conditions.

Our mechanistic studies demonstrate that TSC-null EV promote adherent 621-101 cell migration and invasion. The TSC-null mediate an increase in EV migration associate with increased actin polymerization and activation of paxillin and ITGα6/β1-c-Src-FAK-AKT regulatory axis, which is alleviated by the blockade of EV uptake or inhibition of c-Src. Our data are consistent with alterations in EV cargo of LAM surrogate cells that enhance VEGF secretion and viability of recipient fibroblast compared to TSC2 addback cells-derived EV cargo^10^, and roles of EV-ITG-β1/5 and c-Src in the regulation of cell adhesion and disease progression in human osteosarcoma^112^. More importantly, this work provides evidence for the previously unknow roles of different EV subtypes in mediating tumor progression that are potentially applicable to other malignancies. These different TSC-null EV subtypes enhance CSC and metastable phenotypes of LAM surrogate CSCs to a different magnitude and through different mechanisms. Using 2D and 3D culture system we showed that primary tumor-like EV are more powerful in enhancing accumulated distance and velocity of migrated sphere cells compared to metastasizing tumor-like EV. The upregulation of OXPHOS genes and increases in ATP levels in spheres treated with primary tumor-like TSC-null EV prior to the initiation of migration are likely mediated by increased accumulation of critical mitochondrial function regulator Nrf2^104^ in these EV, and associate with moderate whole-cell activation of AMPK. This is further corroborated by Nrf2 enrichment in LAM-EV compared to Normal-EV. This metabolic and EV mediated shift in LAM cells, associated with enrichment of activated AMPK, TFAM, ATP synthase and increased levels of ATP in chemotactic (FBS) pseudopodia (Pd), indicates novel EV function in coupling local energy demands to subcellular targeting of energy source for the activation of migratory machinery, facilitating faster and more distant cell migration. Our data are consistent with not only AMPK function as essential energy sensor and metabolic regulator^84^, but also with AMPK mediated subcellular targeting of mitochondria to the leading edge and protrusive structures in the response to local energy demands during cell migration and invasion^84^. They are also consistent with previous report indicating that cell protrusions of migrated cells are on high energy demand and that local AMPK activation fulfills these demands^84^. Our data reveal novel and unreported function of primary tumor-like TSC-null EV in regulating plasticity of cell migration via localized AMPK activation and subcellular mitochondria and ATP synthase localization, and support previous notion of heterogeneity of cellular energy balance^84^. In addition, these primary tumor-like EV are also more powerful in enhancing lung seeding by circulated LAM cells *in vivo* compared to metastasizing tumor-like EV, which is consistent with primary tumor-like EV superiority in promoting cell migration. The explanation for these phenotypes is likely in mechanisms utilized by these two types of TSC-null EV. While primary tumor-like EV promoted localized ATP synthesis, and thus, faster migration, the metastasizing tumor-like EV promoted formation of IAC. The IAC are organized into subdomains and functional modules to mediate its assembly^105^. They are also organized vertically into functional layers, including adjacent to the membrane integrin signaling layer, composed of integrins, FAK, paxillin and ILK; intermediated force-transduction layer with talin and vinculin proteins, and actin-regulatory layer enriched with actin-associated proteins^105^. Our data indicate that metastasizing tumor-like TSC-null EV facilitate migration of LAM cell via formation of IAC and IAC signaling through delivery of EV pool of IAC building blocks, including vinculin, paxillin and ILK1, similar to the function of cytosolic pool of these blocks^105^. This is also corroborated by enrichment of paxillin and ILK1 in LAM-EV compared to Normal-EV.

In integrin-dependent migration modes, different velocities come from different level of adhesion strength^113^. The slightly lower velocities and accumulated distance of migrated sphere cells treated with metastasizing vs. primary tumor-like EV might be explained by formation of IAC itself and stronger adhesions. Of note, the superiority of primary tumor-like EV over metastasizing tumor-like EV in promoting cell migration was alleviated when migration was examined using transwell assay and after sphere dissociation. This discrepancy could be explained by dissociation procedures interfering with mitochondrial function and localization, underscoring the necessity of experimental design mimicking *in vivo* conditions. Nonetheless, the metastasizing tumor-like EV were superior in promoting stemness of LAM CSCs indicated by sphere size, ALDH activity, and increased CD44 expression, which is consistent with role of EV of Ewing sarcoma in promoting CSC^114,115^. These data also suggest that the main role of this EV subtype is CSC protection with secondary but significant influence on CSC migration and lung seeding by circulating cells. Both type of EV shared equal contribution to CSC invasion indicating their importance in this process.

Overall our study revealed unknown functions of different EV subtypes that are derived from genetically identical tumor cells and different machinery engagement to facilitate these functions. We also demonstrated that LAM-EV share several features with TSC-null EV and are enriched with various pathways for DEP that play multiple roles in tumor progression, including well established role of estrogen signaling and metabolic alterations in LAM. Therefore, targeting EV pathway for LAM could be potentially an effective therapeutic strategy that is superior to single agent therapy against LAM.

## Acknowledgement

This work has been supported by NIH NHLBI R01HL160972 to M.K and J.Y.

## References

1 Karbowniczek, M. et al. Recurrent lymphangiomyomatosis after transplantation: genetic analyses reveal a metastatic mechanism. Am J Respir Crit Care Med 167, 976–982, doi:10.1164/rccm.200208-969OC (2003).

2 Prizant, H. et al. Uterine-specific loss of Tsc2 leads to myometrial tumors in both the uterus and lungs. Molecular endocrinology 27, 1403–1414, doi:10.1210/me.2013-1059 (2013).

3 Henske, E. P. & McCormack, F. X. Lymphangioleiomyomatosis - a wolf in sheep’s clothing. J Clin Invest 122, 3807–3816, doi:10.1172/JCI58709 (2012).

4 Guo, M. et al. Single-Cell Transcriptomic Analysis Identifies a Unique Pulmonary Lymphangioleiomyomatosis Cell. Am J Respir Crit Care Med 202, 1373–1387, doi:10.1164/rccm.201912-2445OC (2020).

5 Carsillo, T., Astrinidis, A. & Henske, E. P. Mutations in the tuberous sclerosis complex gene TSC2 are a cause of sporadic pulmonary lymphangioleiomyomatosis. Proc. Natl. Acad. Sci. U. S. A. 97, 6085–6090, doi:DOI 10.1073/pnas.97.11.6085 (2000).

6 Astrinidis, A. et al. Mutational analysis of the tuberous sclerosis gene TSC2 in patients with pulmonary lymphangioleiomyomatosis. J. Med. Genet. 37, 55–57, doi:DOI 10.1136/jmg.37.1.55 (2000).

7 Plank, T. L., Yeung, R. S. & Henske, E. P. Hamartin, the product of the tuberous sclerosis 1 (TSC1) gene, interacts with tuberin and appears to be localized to cytoplasmic vesicles. Cancer Res 58, 4766–4770 (1998).

8 Tee, A. R., Manning, B. D., Roux, P. P., Cantley, L. C. & Blenis, J. Tuberous sclerosis complex gene products, Tuberin and Hamartin, control mTOR signaling by acting as a GTPase-activating protein complex toward Rheb. Curr Biol 13, 1259–1268 (2003).

9 Patel, B. et al. Exosomes mediate the acquisition of the disease phenotypes by cells with normal genome in tuberous sclerosis complex. Oncogene 35, 3027–3036, doi:10.1038/onc.2015.358 (2016).

10 Bhaoighill, M. N. et al. Tuberous Sclerosis Complex cell-derived EVs have an altered protein cargo capable of regulating their microenvironment and have potential as disease biomarkers. J Extracell Vesicles 12, e12336, doi:10.1002/jev2.12336 (2023).

11 Li, Y., Yin, Z., Fan, J., Zhang, S. & Yang, W. The roles of exosomal miRNAs and lncRNAs in lung diseases. Signal Transduct Target Ther 4, 47, doi:10.1038/s41392-019-0080-7 (2019).

12 Mohan, A., Agarwal, S., Clauss, M., Britt, N. S. & Dhillon, N. K. Extracellular vesicles: novel communicators in lung diseases. Respir Res 21, 175, doi:10.1186/s12931-020-01423-y (2020).

13 Burgy, O. et al. New players in chronic lung disease identified at the European Respiratory Society International Congress in Paris 2018: from microRNAs to extracellular vesicles. J Thorac Dis 10, S2983–S2987, doi:10.21037/jtd.2018.08.20 (2018).

14 Kubo, H. Extracellular Vesicles in Lung Disease. Chest 153, 210–216, doi:10.1016/j.chest.2017.06.026 (2018).

15 Bastos, N., Ruivo, C. F., da Silva, S. & Melo, S. A. Exosomes in cancer: Use them or target them? Semin Cell Dev Biol 78, 13–21, doi:10.1016/j.semcdb.2017.08.009 (2018).

16 Atai, N. A. et al. Heparin blocks transfer of extracellular vesicles between donor and recipient cells. J Neurooncol 115, 343–351, doi:10.1007/s11060-013-1235-y (2013).

17 Christianson, H. C., Svensson, K. J., van Kuppevelt, T. H., Li, J. P. & Belting, M. Cancer cell exosomes depend on cell-surface heparan sulfate proteoglycans for their internalization and functional activity. Proc Natl Acad Sci U S A 110, 17380–17385, doi:10.1073/pnas.1304266110 (2013).

18 Chen, C. C. et al. Elucidation of Exosome Migration across the Blood-Brain Barrier Model In Vitro. Cell Mol Bioeng 9, 509–529, doi:10.1007/s12195-016-0458-3 (2016).

19 Svensson, K. J. et al. Exosome uptake depends on ERK1/2-heat shock protein 27 signaling and lipid Raft-mediated endocytosis negatively regulated by caveolin-1. J Biol Chem 288, 17713–17724, doi:10.1074/jbc.M112.445403 (2013).

20 McCluskey, A. et al. Building a better dynasore: the dyngo compounds potently inhibit dynamin and endocytosis. Traffic 14, 1272–1289, doi:10.1111/tra.12119 (2013).

21 Hazan-Halevy, I. et al. Cell-specific uptake of mantle cell lymphoma-derived exosomes by malignant and non-malignant B-lymphocytes. Cancer Lett 364, 59–69, doi:10.1016/j.canlet.2015.04.026 (2015).

22 Essandoh, K. et al. Blockade of exosome generation with GW4869 dampens the sepsis-induced inflammation and cardiac dysfunction. Biochim Biophys Acta 1852, 2362–2371, doi:10.1016/j.bbadis.2015.08.010 (2015).

23 Datta, A. et al. High-throughput screening identified selective inhibitors of exosome biogenesis and secretion: A drug repurposing strategy for advanced cancer. Sci Rep 8, 8161, doi:10.1038/s41598-018-26411-7 (2018).

24 Sun, Z., Wang, L., Dong, L. & Wang, X. Emerging role of exosome signalling in maintaining cancer stem cell dynamic equilibrium. J Cell Mol Med, doi:10.1111/jcmm.13676 (2018).

25 Kalluri, R. The biology and function of exosomes in cancer. J Clin Invest 126, 1208–1215, doi:10.1172/JCI81135 (2016).

26 Feng, W., Dean, D. C., Hornicek, F. J., Shi, H. & Duan, Z. Exosomes promote pre-metastatic niche formation in ovarian cancer. Mol Cancer 18, 124, doi:10.1186/s12943-019-1049-4 (2019).

27 Mashouri, L. et al. Exosomes: composition, biogenesis, and mechanisms in cancer metastasis and drug resistance. Mol Cancer 18, 75, doi:10.1186/s12943-019-0991-5 (2019).

28 Hu, L. Z., Wickline, S. A. & Hood, J. L. Magnetic resonance imaging of melanoma exosomes in lymph nodes. Magn Reson Med 74, 266–271, doi:10.1002/mrm.25376 (2015).

29 Kim, J. et al. Replication study: Melanoma exosomes educate bone marrow progenitor cells toward a pro-metastatic phenotype through MET. Elife 7, doi:10.7554/eLife.39944 (2018).

30 Azmi, A. S., Bao, B. & Sarkar, F. H. Exosomes in cancer development, metastasis, and drug resistance: a comprehensive review. Cancer Metast Rev 32, 623–642, doi:10.1007/s10555-013-9441-9 (2013).

31 Peinado, H. et al. Melanoma exosomes educate bone marrow progenitor cells toward a pro-metastatic phenotype through MET. Nature medicine 18, 883–891, doi:10.1038/nm.2753 (2012).

32 Hood, J. L., Roman, S. S. & Wickline, S. A. Exosomes Released by Melanoma Cells Prepare Sentinel Lymph Nodes for Tumor Metastasis. Cancer Research 71, 3792–3801, doi:10.1158/0008-5472.CAN-10-4455 (2011).

33 DeRita, R. M. et al. Tumor-Derived Extracellular Vesicles Require beta1 Integrins to Promote Anchorage-Independent Growth. iScience 14, 199–209, doi:10.1016/j.isci.2019.03.022 (2019).

34 DeRita, R. M. et al. c-Src, Insulin-Like Growth Factor I Receptor, G-Protein-Coupled Receptor Kinases and Focal Adhesion Kinase are Enriched Into Prostate Cancer Cell Exosomes. J Cell Biochem 118, 66–73, doi:10.1002/jcb.25611 (2017).

35 Hoshino, A. et al. Tumour exosome integrins determine organotropic metastasis. Nature 527, 329–335, doi:10.1038/nature15756 (2015).

36 Grum-Schwensen, B. et al. Suppression of tumor development and metastasis formation in mice lacking the S100A4(mts1) gene. Cancer Res 65, 3772–3780, doi:10.1158/0008-5472.CAN-04-4510 (2005).

37 Lukanidin, E. & Sleeman, J. P. Building the niche: the role of the S100 proteins in metastatic growth. Semin Cancer Biol 22, 216–225, doi:10.1016/j.semcancer.2012.02.006 (2012).

38 Liu, L. et al. S100A4 alters metabolism and promotes invasion of lung cancer cells by up-regulating mitochondrial complex I protein NDUFS2. J Biol Chem, doi:10.1074/jbc.RA118.004365 (2019).

39 Semov, A. et al. Metastasis-associated protein S100A4 induces angiogenesis through interaction with Annexin II and accelerated plasmin formation. J Biol Chem 280, 20833–20841, doi:10.1074/jbc.M412653200 (2005).

40 Jia, W., Gao, X. J., Zhang, Z. D., Yang, Z. X. & Zhang, G. S100A4 silencing suppresses proliferation, angiogenesis and invasion of thyroid cancer cells through downregulation of MMP-9 and VEGF. Eur Rev Med Pharmacol Sci 17, 1495–1508 (2013).

41 Hernandez, J. L. et al. Therapeutic targeting of tumor growth and angiogenesis with a novel anti-S100A4 monoclonal antibody. PLoS One 8, e72480, doi:10.1371/journal.pone.0072480 (2013).

42 Masaoutis, C., Korkolopoulou, P. & Theocharis, S. Exosomes in sarcomas: Tiny messengers with broad implications in diagnosis, surveillance, prognosis and treatment. Cancer Lett 449, 172–177, doi:10.1016/j.canlet.2019.02.025 (2019).

43 Min, L., Shen, J., Tu, C., Hornicek, F. & Duan, Z. The roles and implications of exosomes in sarcoma. Cancer Metastasis Rev 35, 377–390, doi:10.1007/s10555-016-9630-4 (2016).

44 Chicon-Bosch, M. & Tirado, O. M. Exosomes in Bone Sarcomas: Key Players in Metastasis. Cells 9, doi:10.3390/cells9010241 (2020).

45 Jolly, M. K., Ware, K. E., Gilja, S., Somarelli, J. A. & Levine, H. EMT and MET: necessary or permissive for metastasis? Mol Oncol 11, 755–769, doi:10.1002/1878-0261.12083 (2017).

46 Granados, K., Poelchen, J., Novak, D. & Utikal, J. Cellular Reprogramming-A Model for Melanoma Cellular Plasticity. Int J Mol Sci 21, doi:10.3390/ijms21218274 (2020).

47 Conigliaro, A. & Cicchini, C. Exosome-Mediated Signaling in Epithelial to Mesenchymal Transition and Tumor Progression. J Clin Med 8, doi:10.3390/jcm8010026 (2018).

48 Yu, M. et al. Circulating breast tumor cells exhibit dynamic changes in epithelial and mesenchymal composition. Science 339, 580–584, doi:10.1126/science.1228522 (2013).

49 Huang, R. Y. et al. An EMT spectrum defines an anoikis-resistant and spheroidogenic intermediate mesenchymal state that is sensitive to e-cadherin restoration by a src-kinase inhibitor, saracatinib (AZD0530). Cell Death Dis 4, e915, doi:10.1038/cddis.2013.442 (2013).

50 Schliekelman, M. J. et al. Molecular portraits of epithelial, mesenchymal, and hybrid States in lung adenocarcinoma and their relevance to survival. Cancer Res 75, 1789–1800, doi:10.1158/0008-5472.CAN-14-2535 (2015).

51 Pastushenko, I. et al. Identification of the tumour transition states occurring during EMT. Nature 556, 463–468, doi:10.1038/s41586-018-0040-3 (2018).

52 Ruscetti, M., Quach, B., Dadashian, E. L., Mulholland, D. J. & Wu, H. Tracking and Functional Characterization of Epithelial-Mesenchymal Transition and Mesenchymal Tumor Cells during Prostate Cancer Metastasis. Cancer Res 75, 2749–2759, doi:10.1158/0008-5472.CAN-14-3476 (2015).

53 Yamashita, N. et al. Epithelial Paradox: Clinical Significance of Coexpression of E-cadherin and Vimentin With Regard to Invasion and Metastasis of Breast Cancer. Clin Breast Cancer 18, e1003–e1009, doi:10.1016/j.clbc.2018.02.002 (2018).

54 Conigliaro, A. et al. Evidence for a common progenitor of epithelial and mesenchymal components of the liver. Cell Death Differ 20, 1116–1123, doi:10.1038/cdd.2013.49 (2013).

55 Sannino, G., Marchetto, A., Kirchner, T. & Grunewald, T. G. P. Epithelial-to-Mesenchymal and Mesenchymal-to-Epithelial Transition in Mesenchymal Tumors: A Paradox in Sarcomas? Cancer Res 77, 4556–4561, doi:10.1158/0008-5472.CAN-17-0032 (2017).

56 Kahlert, U. D., Joseph, J. V. & Kruyt, F. A. E. EMT- and MET-related processes in nonepithelial tumors: importance for disease progression, prognosis, and therapeutic opportunities. Mol Oncol 11, 860–877, doi:10.1002/1878-0261.12085 (2017).

57 Yang, J. et al. Mesenchymal to epithelial transition in sarcomas. Eur J Cancer 50, 593–601, doi:10.1016/j.ejca.2013.11.006 (2014).

58 Somarelli, J. A. et al. Mesenchymal-Epithelial Transition in Sarcomas Is Controlled by the Combinatorial Expression of MicroRNA 200s and GRHL2. Mol Cell Biol 36, 2503–2513, doi:10.1128/MCB.00373-16 (2016).

59 Qi, Y. et al. Transforming growth factor-beta1 signaling promotes epithelial-mesenchymal transition-like phenomena, cell motility, and cell invasion in synovial sarcoma cells. PLoS One 12, e0182680, doi:10.1371/journal.pone.0182680 (2017).

60 Martinez-Delgado, P. et al. Cancer Stem Cells in Soft-Tissue Sarcomas. Cells 9, doi:10.3390/cells9061449 (2020).

61 Genadry, K. C., Pietrobono, S., Rota, R. & Linardic, C. M. Soft Tissue Sarcoma Cancer Stem Cells: An Overview. Front Oncol 8, 475, doi:10.3389/fonc.2018.00475 (2018).

62 Seyama, K., Kumasaka, T., Kurihara, M., Mitani, K. & Sato, T. Lymphangioleiomyomatosis: a disease involving the lymphatic system. Lymphat Res Biol 8, 21–31, doi:10.1089/lrb.2009.0018 (2010).

63 Cai, X. et al. Phenotypic characterization of disseminated cells with TSC2 loss of heterozygosity in patients with lymphangioleiomyomatosis. Am J Respir Crit Care Med 182, 1410–1418, doi:10.1164/rccm.201003-0489OC (2010).

64 Pacheco-Rodriguez, G. et al. Circulating Lymphangioleiomyomatosis Tumor Cells With Loss of Heterozygosity in the TSC2 Gene Show Increased Aldehyde Dehydrogenase Activity. Chest 156, 298–307, doi:10.1016/j.chest.2019.03.040 (2019).

65 Grzegorek, I. et al. Immunohistochemical evaluation of pulmonary lymphangioleiomyomatosis. Anticancer Res 35, 3353–3360 (2015).

66 Tang, Y., Kwiatkowski, D. J. & Henske, E. P. Midkine expression by stem-like tumor cells drives persistence to mTOR inhibition and an immune-suppressive microenvironment. Nat Commun 13, 5018, doi:10.1038/s41467-022-32673-7 (2022).

67 Howe, S. R., Gottardis, M. M., Everitt, J. I. & Walker, C. Estrogen stimulation and tamoxifen inhibition of leiomyoma cell growth in vitro and in vivo. Endocrinology 136, 4996–5003, doi:10.1210/endo.136.11.7588234 (1995).

68 Astrinidis, A. et al. Tuberin, the tuberous sclerosis complex 2 tumor suppressor gene product, regulates Rho activation, cell adhesion and migration. Oncogene 21, 8470–8476, doi:10.1038/sj.onc.1205962 (2002).

69 Yu, J., Astrinidis, A., Howard, S. & Henske, E. P. Estradiol and tamoxifen stimulate LAM-associated angiomyolipoma cell growth and activate both genomic and nongenomic signaling pathways. Am J Physiol Lung Cell Mol Physiol 286, L694–700 (2004).

70 Hong, F. et al. mTOR-raptor binds and activates SGK1 to regulate p27 phosphorylation. Molecular cell 30, 701–711, doi:10.1016/j.molcel.2008.04.027 (2008).

71 Astrinidis, A. et al. Upregulation of acid ceramidase contributes to tumor progression in tuberous sclerosis complex. JCI Insight 8, doi:10.1172/jci.insight.166850 (2023).

72 Thery, C. et al. Minimal information for studies of extracellular vesicles 2018 (MISEV2018): a position statement of the International Society for Extracellular Vesicles and update of the MISEV2014 guidelines. J Extracell Vesicles 7, 1535750, doi:10.1080/20013078.2018.1535750 (2018).

73 Wang, Y. J., Bailey, J. M., Rovira, M. & Leach, S. D. Sphere-forming assays for assessment of benign and malignant pancreatic stem cells. Methods Mol Biol 980, 281–290, doi:10.1007/978-1-62703-287-2_15 (2013).

74 Cho, J. H. et al. Notch transactivates Rheb to maintain the multipotency of TSC-null cells. Nat Commun 8, 1848, doi:10.1038/s41467-017-01845-1 (2017).

75 Sung, B. H. et al. A live cell reporter of exosome secretion and uptake reveals pathfinding behavior of migrating cells. Nat Commun 11, 2092, doi:10.1038/s41467-020-15747-2 (2020).

76 Matoba, Y. et al. Targeting Galectin 3 illuminates its contributions to the pathology of uterine serous carcinoma. Br J Cancer 130, 1463–1476, doi:10.1038/s41416-024-02621-x (2024).

77 Kowal, J. et al. Proteomic comparison defines novel markers to characterize heterogeneous populations of extracellular vesicle subtypes. Proc Natl Acad Sci U S A 113, E968–977, doi:10.1073/pnas.1521230113 (2016).

78 Gupta, S. et al. An improvised one-step sucrose cushion ultracentrifugation method for exosome isolation from culture supernatants of mesenchymal stem cells. Stem Cell Res Ther 9, 180, doi:10.1186/s13287-018-0923-0 (2018).

79 Colombo, M., Raposo, G. & Thery, C. Biogenesis, secretion, and intercellular interactions of exosomes and other extracellular vesicles. Annu Rev Cell Dev Biol 30, 255–289, doi:10.1146/annurev-cellbio-101512-122326 (2014).

80 Lasser, C., Eldh, M. & Lotvall, J. Isolation and characterization of RNA-containing exosomes. J Vis Exp, e3037, doi:10.3791/3037 (2012).

81 Rikkert, L. G., Nieuwland, R., Terstappen, L. & Coumans, F. A. W. Quality of extracellular vesicle images by transmission electron microscopy is operator and protocol dependent. J Extracell Vesicles 8, 1555419, doi:10.1080/20013078.2018.1555419 (2019).

82 Otomo, M. et al. Some selective serotonin reuptake inhibitors inhibit dynamin I guanosine triphosphatase (GTPase). Biol Pharm Bull 31, 1489–1495 (2008).

83 Dong, G. et al. DDX18 drives tumor immune escape through transcription-activated STAT1 expression in pancreatic cancer. Oncogene 42, 3000–3014, doi:10.1038/s41388-023-02817-0 (2023).

84 Cunniff, B., McKenzie, A. J., Heintz, N. H. & Howe, A. K. AMPK activity regulates trafficking of mitochondria to the leading edge during cell migration and matrix invasion. Mol Biol Cell 27, 2662–2674, doi:10.1091/mbc.E16-05-0286 (2016).

85 Jones, M. C. et al. Isolation of integrin-based adhesion complexes. Curr Protoc Cell Biol 66, 9 8 1–9 8 15, doi:10.1002/0471143030.cb0908s66 (2015).

86 Langmead, B. & Salzberg, S. L. Fast gapped-read alignment with Bowtie 2. Nat Methods 9, 357–359, doi:10.1038/nmeth.1923 (2012).

87 Li, B. & Dewey, C. N. RSEM: accurate transcript quantification from RNA-Seq data with or without a reference genome. BMC Bioinformatics 12, 323, doi:10.1186/1471-2105-12-323 (2011).

88 Robinson, M. D., McCarthy, D. J. & Smyth, G. K. edgeR: a Bioconductor package for differential expression analysis of digital gene expression data. Bioinformatics 26, 139–140, doi:10.1093/bioinformatics/btp616 (2010).

89 Young, M. D., Wakefield, M. J., Smyth, G. K. & Oshlack, A. Gene ontology analysis for RNA-seq: accounting for selection bias. Genome Biol 11, R14, doi:10.1186/gb-2010-11-2-r14 (2010).

90 Xie, C. et al. KOBAS 2.0: a web server for annotation and identification of enriched pathways and diseases. Nucleic Acids Res 39, W316–322, doi:10.1093/nar/gkr483 (2011).

91 Parkhomchuk, D. et al. Transcriptome analysis by strand-specific sequencing of complementary DNA. Nucleic Acids Res 37, e123, doi:10.1093/nar/gkp596 (2009).

92 Mortazavi, A., Williams, B. A., McCue, K., Schaeffer, L. & Wold, B. Mapping and quantifying mammalian transcriptomes by RNA-Seq. Nat Methods 5, 621–628, doi:10.1038/nmeth.1226 (2008).

93 Liao, Y., Smyth, G. K. & Shi, W. featureCounts: an efficient general purpose program for assigning sequence reads to genomic features. Bioinformatics 30, 923–930, doi:10.1093/bioinformatics/btt656 (2014).

94 Love, M. I., Huber, W. & Anders, S. Moderated estimation of fold change and dispersion for RNA-seq data with DESeq2. Genome Biol 15, 550, doi:10.1186/s13059-014-0550-8 (2014).

95 Anders, S. & Huber, W. Differential expression analysis for sequence count data. Genome Biol 11, R106, doi:10.1186/gb-2010-11-10-r106 (2010).

96 Kanehisa, M. & Goto, S. KEGG: kyoto encyclopedia of genes and genomes. Nucleic Acids Res 28, 27–30, doi:10.1093/nar/28.1.27 (2000).

97 Walker, J. A. et al. Quantitative PCR for DNA identification based on genome-specific interspersed repetitive elements. Genomics 83, 518–527, doi:10.1016/j.ygeno.2003.09.003 (2004).

98 Yu, J. J. et al. Estrogen promotes the survival and pulmonary metastasis of tuberin-null cells. Proc Natl Acad Sci U S A 106, 2635–2640, doi:10.1073/pnas.0810790106 (2009).

99 Hayashi, T. et al. Immunohistochemical study of matrix metalloproteinases (MMPs) and their tissue inhibitors (TIMPs) in pulmonary lymphangioleiomyomatosis (LAM). Hum Pathol 28, 1071–1078, doi:10.1016/s0046-8177(97)90061-7 (1997).

100 Matsui, K. et al. Role for activation of matrix metalloproteinases in the pathogenesis of pulmonary lymphangioleiomyomatosis. Arch Pathol Lab Med 124, 267–275, doi:10.5858/2000-124-0267-RFAOMM (2000).

101 Odajima, N. et al. Matrix metalloproteinases in blood from patients with LAM. Respir Med 103, 124–129, doi:10.1016/j.rmed.2008.07.017 (2009).

102 Tyryshkin, A., Bhattacharya, A. & Eissa, N. T. SRC kinase is a novel therapeutic target in lymphangioleiomyomatosis. Cancer Res 74, 1996–2005, doi:10.1158/0008-5472.CAN-13-1256 (2014).

103 Pacheco-Rodriguez, G. et al. TSC2 loss in lymphangioleiomyomatosis cells correlated with expression of CD44v6, a molecular determinant of metastasis. Cancer Res 67, 10573–10581, doi:10.1158/0008-5472.CAN-07-1356 (2007).

104 Dinkova-Kostova, A. T. & Abramov, A. Y. The emerging role of Nrf2 in mitochondrial function. Free Radic Biol Med 88, 179–188, doi:10.1016/j.freeradbiomed.2015.04.036 (2015).

105 Chastney, M. R., Conway, J. R. W. & Ivaska, J. Integrin adhesion complexes. Curr Biol 31, R536–R542, doi:10.1016/j.cub.2021.01.038 (2021).

106 Horton, E. R. et al. Modulation of FAK and Src adhesion signaling occurs independently of adhesion complex composition. J Cell Biol 212, 349–364, doi:10.1083/jcb.201508080 (2016).

107 Sun, Y. et al. Estradiol promotes pentose phosphate pathway addiction and cell survival via reactivation of Akt in mTORC1 hyperactive cells. Cell Death Dis 5, e1231, doi:10.1038/cddis.2014.204 (2014).

108 Hakulinen, J., Sankkila, L., Sugiyama, N., Lehti, K. & Keski-Oja, J. Secretion of active membrane type 1 matrix metalloproteinase (MMP-14) into extracellular space in microvesicular exosomes. J Cell Biochem 105, 1211–1218, doi:10.1002/jcb.21923 (2008).

109 Zadjali, F. et al. Tuberous Sclerosis Complex Axis Controls Renal Extracellular Vesicle Production and Protein Content. Int J Mol Sci 21, doi:10.3390/ijms21051729 (2020).

110 Kumar, P. et al. Tsc Gene Locus Disruption and Differences in Renal Epithelial Extracellular Vesicles. Front Physiol 12, 630933, doi:10.3389/fphys.2021.630933 (2021).

111 Zou, W. et al. Exosome Release Is Regulated by mTORC1. Adv Sci (Weinh) 6, 1801313, doi:10.1002/advs.201801313 (2019).

112 Jerez, S. et al. Proteomic Analysis of Exosomes and Exosome-Free Conditioned Media From Human Osteosarcoma Cell Lines Reveals Secretion of Proteins Related to Tumor Progression. J Cell Biochem 118, 351–360, doi:10.1002/jcb.25642 (2017).

113 Schmidt, S. & Friedl, P. Interstitial cell migration: integrin-dependent and alternative adhesion mechanisms. Cell Tissue Res 339, 83–92, doi:10.1007/s00441-009-0892-9 (2010).

114 Kling, M. J. et al. Exosomes secreted under hypoxia enhance stemness in Ewing’s sarcoma through miR-210 delivery. Oncotarget 11, 3633–3645, doi:10.18632/oncotarget.27702 (2020).

115 De Feo, A. et al. Exosomes from CD99-deprived Ewing sarcoma cells reverse tumor malignancy by inhibiting cell migration and promoting neural differentiation. Cell Death Dis 10, 471, doi:10.1038/s41419-019-1675-1 (2019).

