## Supplementary Figures for "Extracellular vesicles regulate metastable phenotypes of lymphangioleiomyomatosis cells via shuttling ATP synthesis to pseudopodia and activation of integrin adhesion complexes"

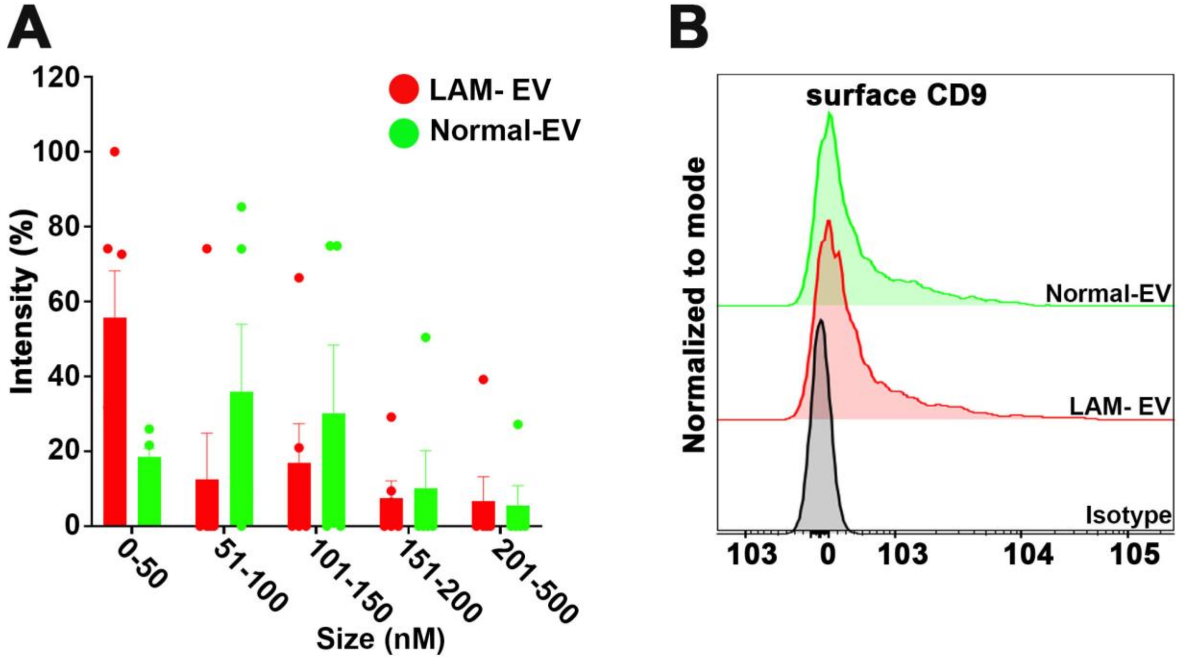


**Supplementary Figure 1. Characterization of LAM-EV & Normal-EV.** (A) EV size distribution by DLS. (B). EV Surface CD9 by FACS. Each Peak is normalized to its mode.

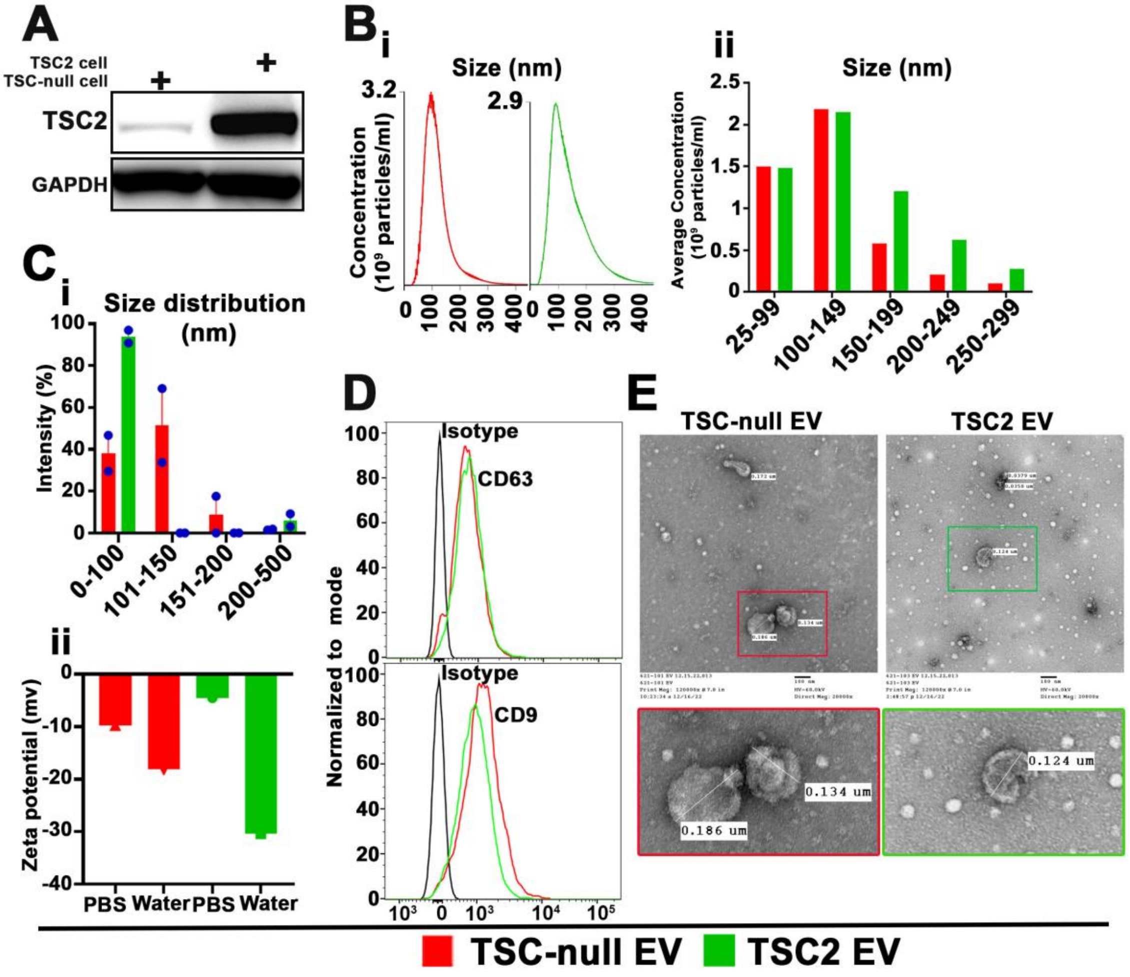


**Supplementary Figure 2. Loss of TSC1/2 alters biochemical and physical characteristics of EV.** (A) Representative immunoblot of TSC2 expression in TSC-null and TSC2 addback cell. (B) NTA analysis of EV derived from TSC-null and TSC2 addback cells, (B-i) particle concentration, (B-ii) size distribution. (C) EV size distribution by DLS analysis. (D) Surface expression of CD63 and CD9 EV marker by FACS. Each Peak is normalized to its mode. (E) Morphology of EV by transmission electron microscopy (TEM).

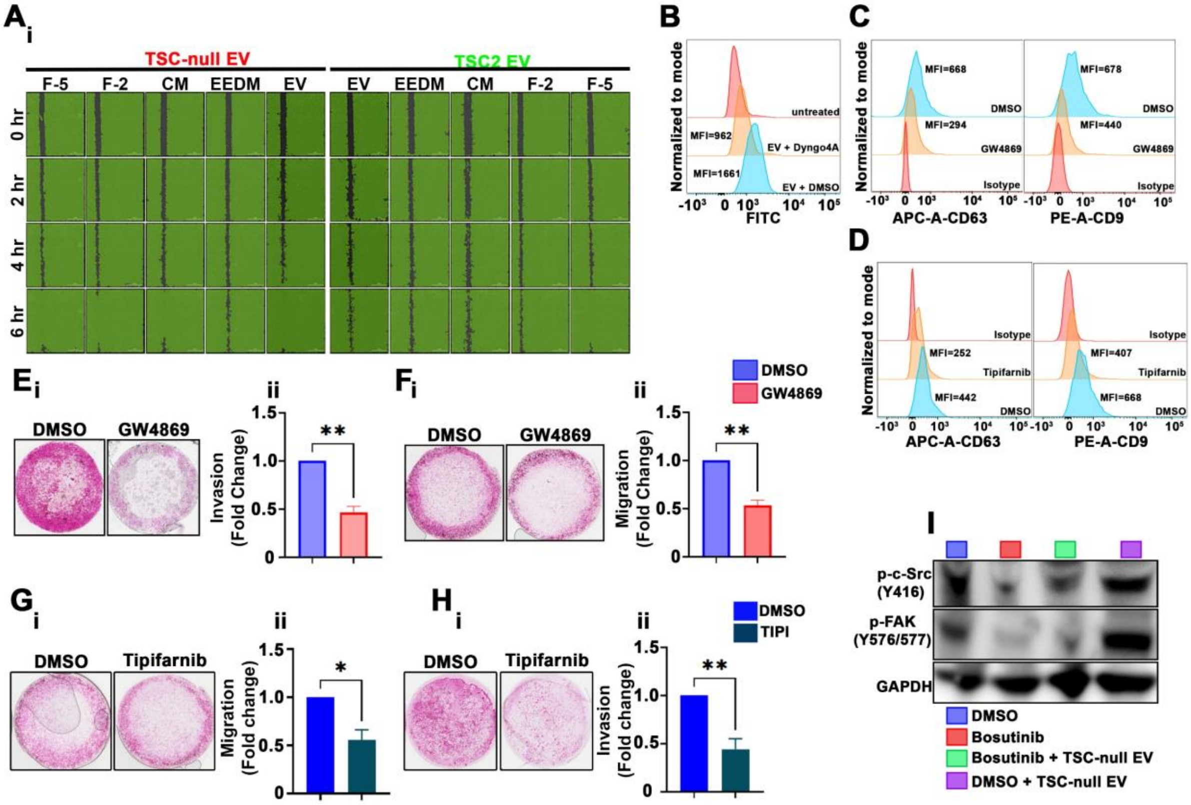


**Supplementary Figure 3. The blockade of TSC-null EV uptake or biogenesis inhibits adherent 621-101 cells migration and invasion.** (A) 621-101 cells were seeded and treated with EV or *JEV* controls and when approximately 95% confluent scratched; (A-i) Representative mark-up images of migrated cells. (A-ii) Quantification of wound closure at indicated timepoint with respect to the wound area at time 0. (B) 621-101 cells were treated with vehicle or EV uptake inhibitor, Dyngo4a, prior adding labelled TSC-null EV. The EV uptake was measured by FACS. Each Peak is normalized to its mode. (C-D) 621-101 cells were treated with vehicle or inhibitor of EV biogenesis, (C) GW4869 or (D) Tipifarnib, and isolated EV were quantified by FACS. Each Peak is normalized to its mode. Histogram shows decrease in EV fluorescence upon (C) GW4869 or (D) Tipifarnib treatment. (E-H) Representative images of the transwell invasion (E-i, G-i,) or migration assay (F-i, H-i), (E-H-ii) quantification of E-H-i, as a fold change in number of cells relative to DMSO. (I) Immunoblot of phospho-c-Src and phopsho-FAK in 621-101 cells treated with Bosutinib. Data are representative of two (B-D) and three (E-I) independent experiments. Bar show mean ± SEM. *P<0.01 by unpaired t-test.

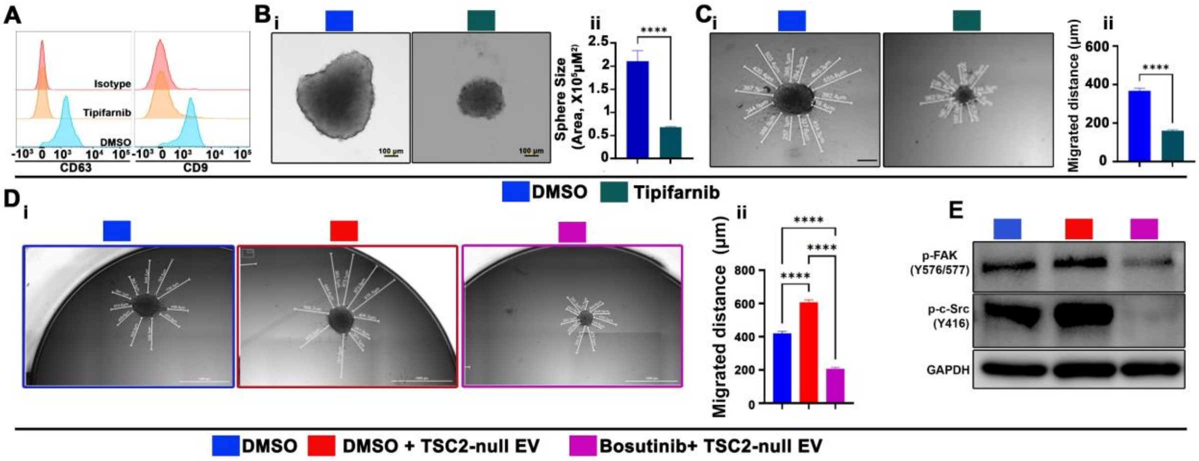


**Supplementary Figure 4. The blockade of TSC-null EV biogenesis inhibits 621-101 sphere size and sphere cell migration.** (A-C) 621-101 spheres were treated with vehicle or inhibitor of EV biogenesis, Tipifarnib (0.1 μM). (A) Isolated EV were quantified by FACS showing surface expression of CD63 and CD9. Histogram shows decrease in EV fluorescence upon Tipifarnib treatment. (B) Sphere assay in 621-101 cells. (B-i) Representative bright field images and (B-ii) quantification of sphere sizes. (C) 621-101 sphere cell migration; (C-i) Representative bright field images and (C-ii) quantification of migrated sphere cells. (D) 621-101 sphere cell migration. Spheres were treated with vehicle or c-Src inhibitor, Bosutinib (1 μM) in the presence or absence of TSC-null EV; (D-i) Representative bright field images and (D-ii) quantification of migrated sphere cells. (E) Phospho-c-Src and phosphor-FAK immunoblot of 621-101 spheres treated with Bosutinib (1 μM). Data represent as Mean ± SEM (n=3). Statistical significance was determined using t-test (B & C) and one-way ANOVA test (D). ****p < 0.0001.

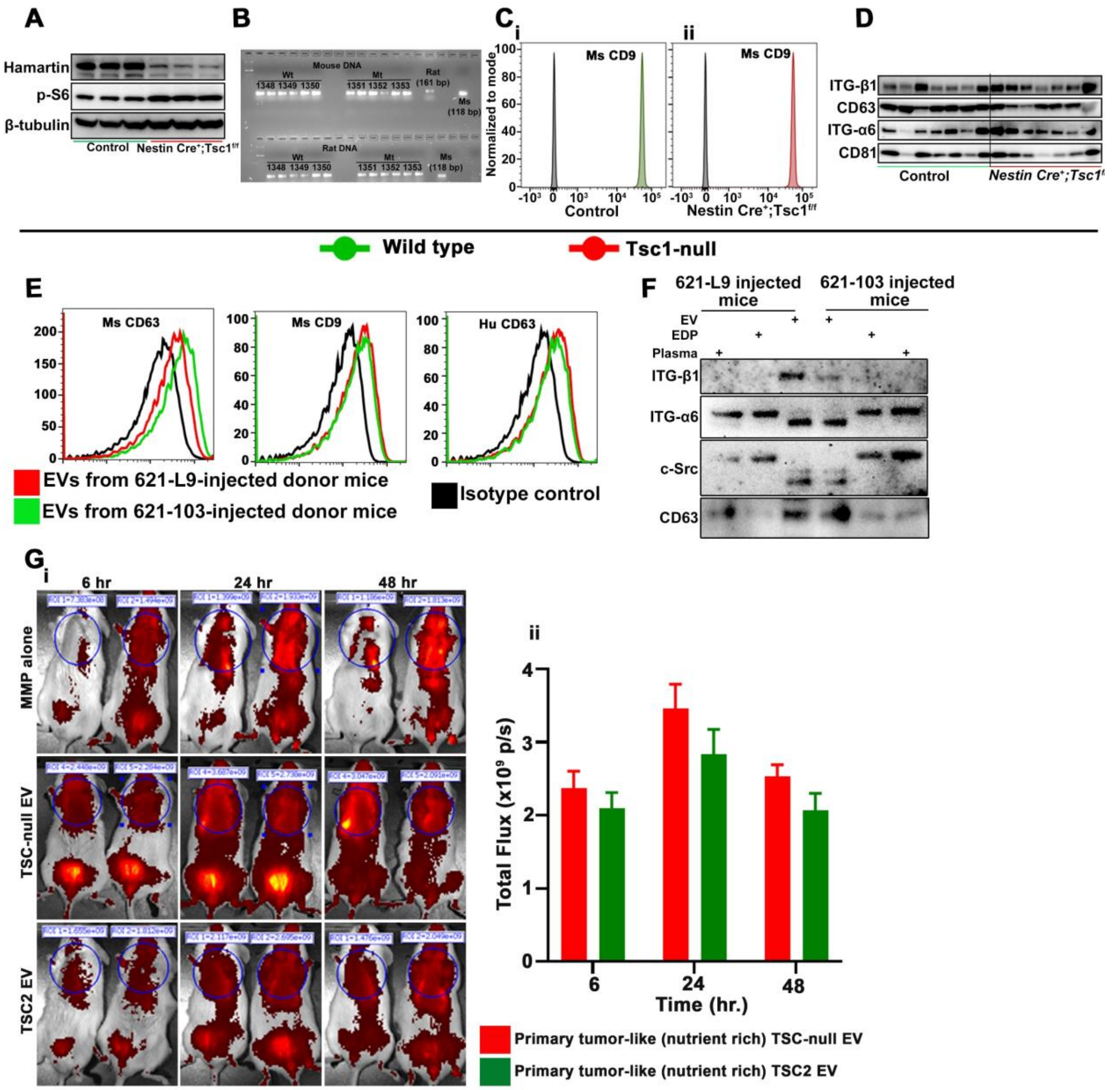


**Supplementary Figure 5.** (A) The expression of hamartin and phospho-S6 in *Tsc1*-null or wild type E15.5 mouse embryo neuronal progenitors' cells by immunoblot. (B) validation of qPCR assays. Amplicons from assays were chromatograph on agarose gel containing Sybr Green Safe DNA gel stain and visualized using UV fluorescence to confirm amplicon size for each primer pair. (C-D) Characterization of EV isolated from cells described in (A) by (C) FACS indicating surface expression of CD9 and (D) immunoblot. (E-F) Characterization of plasma EV isolated from mice injected with 621-L9 or TSC2 addback cells by (E) FACS for surface expression of CD63 and CD9, and (F) immunoblot. (G) Lung MMP activity using IVISSense MMP 750 FAST fluorescent probe from mice treated with primary tumor-like EV i.v. injection 24 hours prior to probe injection. (G-i) Representative fluorescent images at 6, 24, and 48 hours post-MMP probe injection, and (G-ii) quantification of fluorescence photon flux in the chest region (n=4-5/group).
